# Epigenetic cross-talk between Sirt1 and Dnmt1 promotes axonal regeneration after spinal cord injury in zebrafish

**DOI:** 10.1101/2024.04.22.590635

**Authors:** Samudra Gupta, Subhra Prakash Hui

## Abstract

Though spinal cord injury (SCI) causes irreversible sensory and motor impairments in human, adult zebrafish retain the potent regenerative capacity by injury-induced proliferation of central nervous system (CNS)-resident progenitor cells to develop new functional neurons at the lesion site. The hallmark of SCI in zebrafish lies in a series of changes in the epigenetic landscape, specifically DNA methylation and histone modifications. Decoding the post-SCI epigenetic modifications is therefore critical for the development of therapeutic remedies that boost SCI recovery process. Here, we have studied on Sirtuin1 (Sirt1), a non-classical histone deacetylase that potentially play a critical role in neural progenitor cells (NPCs) proliferation and axonal regrowth following SCI in zebrafish. We investigated the role of Sirt1 in NPC proliferation and axonal regrowth in response to injury in the regenerating spinal cord and found that Sirt1 is involved in the induction of NPC proliferation along with glial bridging during spinal cord regeneration. We also demonstrate that Sirt1 plays a pivotal role in regulating the HIPPO pathway through deacetylation-mediated inactivation of Dnmt1 and subsequent hypomethylation of *yap1* promoter, leading to the induction of *ctgfa* expression, which drives the NPC proliferation and axonal regrowth to complete the regenerative process. In conclusion, our study reveals a novel cross-talk between two important epigenetic effectors, Sirt1 and Dnmt1, in the context of spinal cord regeneration, establishing a previously undisclosed relation between Sirt1 and Yap1 which provides a deeper understanding of the underlying mechanisms governing injury-induced NPC proliferation and axonal regrowth. Therefore, we have identified Sirt1 as a novel, major epigenetic regulator of spinal cord regeneration by modulating the HIPPO pathway in zebrafish.

## 1. Introduction

Spinal cord injury (SCI) results from external factors directly or indirectly impacting the spinal cord, often causing partial or complete loss of sensory and motor functions below the injury site (1,2). Epidemiological studies indicate a rising incidence of SCI, posing a significant burden due to its occurrence primarily in young adults and the associated high disability rate (3). Enhancing the structural and functional reconstruction of the injured spinal cord is crucial for improving the quality of life for affected individuals. Despite ongoing efforts, SCI treatment remains a formidable global challenge. Current approaches encompass anti-inflammatory drugs, neuroprotective factors, cell transplantation, and tissue engineering; however, these treatments exhibit limited efficacy in promoting nerve regeneration and functional recovery (4,5). Therefore, investigating the mechanisms of axonal regeneration following neuronal injury is imperative, offering a potential new theoretical foundation for advancing SCI treatment.

Differential responses among glial, neuronal, and systemic components contribute to varying regenerative capacities observed in central nervous system (CNS) tissues across vertebrates (6,7). While SCI in mammals results in irreversible sensory and motor deficits, adult zebrafish exhibit remarkable regenerative potential, achieving functional recovery from paralytic condition within 8 weeks following complete spinal cord transection(8,9). In mammals, astrocytes demonstrate diverse injury responses, often overshadowed by scar-forming cells and inhibitory extracellular molecules that impede regeneration(10). In zebrafish following SCI, specialized glial cells, which are the neural progenitor cells (NPCs),also known as Ependymo-Radial Glial cells (ERG), form a bridge which served as both a physical and signalling scaffold, facilitating axonal regrowth across the lesion(7). Despite extensive investigations into glial cell responses in mammals, our comprehension of how glial cells react to SCI in highly regenerative vertebrates, such as zebrafish, remains limited.

The intricate regulation of stage-specific gene expression in the nervous system during repair and regeneration involves not only transcriptional mechanisms but also encompasses epigenetic modulation. Recent investigations have highlighted that SCI can instigate epigenetic modifications capable of influencing regeneration-associated genes (RAGs), thereby exerting control over the capacity for regeneration and axonal regrowth (11,12).

To explicate the involvement of epigenetic modifiers in the context of spinal cord regeneration, our investigation has centred on Sirtuin 1 (Sirt1), an extensively conserved Nicotinamide Adenine dinucleotide (NAD^+^)–dependent class III protein deacetylase. This enzyme holds potential significance in the regulation of diverse metabolic and pathophysiological phenomena, such as inflammation and aging (13–16). A recent report demonstrated the significant involvement of SIRT1 in the self-renewal and proliferation of mouse embryonic stem cells, particularly under oxidative stress(17). Many other studies have highlighted the neuroprotective role of SIRT1, showing its ability to prevent neuronal damage and enhance long-term neurological function(18,19). Chen et al. reported that SIRT1 promotes recovery of neuronal function and survival by influencing macrophage polarization following spinal cord injury in mice (20). The factor is involved in various neuronal processes, such as regulating differentiation (21,22), providing protection during ischemic preconditioning (23), and preventing neurodegeneration in Alzheimer’s disease(24). In addition, few studies emphasize the essential role of SIRT1 in maintaining normal cognitive function and synaptic plasticity (25,26). However, the precise molecular mechanism through which Sirt1 influences progenitor cell proliferation during spinal cord regeneration in zebrafish remains undefined.

In the present study, we investigated the roles of Sirt1 in NPC proliferation, and as a deacetylase in HIPPO pathway in the context of NPC proliferation in response to injury. We identify Sirt1 as a novel, major regulator of NPC proliferation and axonal regrowth in response to injury in zebrafish spinal cord. We demonstrated modulation of methylation state of Yap1, a major controlling factor in HIPPO pathway, and also elucidate the mechanism by which Sirt1 targets members of the Dnmt family, regulating their deacetylation, and their subsequent role in Yap1 activation and molecular events leading to *ctgfa* expression, which drives Ctgfa-mediated NPC proliferation, glial bridging and subsequent axonal regrowth.

## 2. Materials and Methods

### 2.1. Experimental model and subject details

Adult zebrafish of the AB strain with ages between 4-6 months were used in this study at approximately equal sex ratios. Tg(*ctgfa*:EGFP) zebrafish strain (a generous gift from Dr. C. Patra Laboratory at Agharkar Research Institute, India) was analysed as hemizygote. Fish were housed at approximately 5 fish per litre in custom made aquarium racks and fed three times daily. Water temperature was maintained at 28°C. All zebrafish husbandry and experiments were performed in accordance with institutional and national animal ethics guidelines and approved by the University Committee for the Purpose of Control and Supervision of Experiments on Animals (CPCSEA).

### 2.2. Injury procedures

Adult fishes were anaesthetized for 5 min in 0.02% tricaine methanesulfonate (MS222; Sigma-Aldrich Corporation, St. Louis, MO) before performing any surgical procedures. Spinal cord injury was performed as described in (27). Briefly, a skin incision was made laterally at the level of the dorsal fin to expose the vertebral column, which corresponds to the 15–16th vertebrae as confirmed previously (28). The spinal cord was completely transected with scissors. A fine suture was used to close the muscle and cutaneous wound and fishes were immediately transferred to aquarium water and allowed to regenerate at 28°C. Regenerating spinal cord tissues (from 5-8 samples) at different time points viz. 3 day post injury (dpi), 7 dpi, 10 dpi, 15 dpi and 30 dpi were collected after terminal anaesthesia was applied to the fishes and only spinal cord tissues were dissected out by opening the bony encasing. About 2 mm-length of spinal cord including the regenerating tissue with adjacent uninjured tissue on either side was excised from the whole spinal cord. Sham injury was performed for swimming behaviour analysis as described above without spinal cord transection.

### 2.3. Inhibitor application

1mg of Ex-527 (Sigma-aldrich, Houston, TX, USA) was dissolved in 50 µL of dimethyl sulfoxide (DMSO), and 1 μl of the solution was administered via intra-spinal injection to each fish immediately following spinal cord transection, utilizing a Hamilton syringe (Hamilton Robotics, NV, USA). The fish subjected to this injection procedure were then reintroduced into the aquatic environment. This protocol was systematically repeated over the following four days (up to 5 dpi). In the rescue experiment, Resveratrol (Sigma-Aldrich, Houston, TX, USA), a specific Sirt1 inducer (2 mg/50 µL), was injected at a dosage of 1 µL per fish intra-spinally after the application of Ex-527 for the subsequent four days. A similar procedure was employed to administer 0.1% DMSO as a vehicle for the control group of fish.

### 2.4. Morpholino application

*sirt1* antisense vivo-morpholino (5′-TTTATTTTCGCCGTCCGCCATCTTC-3′) and a standard control morpholino (obtained from Gene Tools, Philomath, USA) were soaked onto small sections of Gelfoam (Pfizer Inc., NY, USA) at a volume of 6 μl each. These Gelfoam pieces were then divided into 30 smaller sections to yield morpholino quantities ranging from 100-800 ng per piece, which were subsequently allowed to dry. For each fish, one of these prepared pieces was applied directly to the site of transection immediately after the procedure. The wound was then sealed using Histoacryl gel (B. Braun, USA), and the animals were monitored for survival over various experimental periods.

### 2.5. Swim path tracking

Zebrafish from various experimental groups, including the control groups, EX527-treated groups and Resveratrol-treated groups, were shifted and acclimatized for 5 minutes to an opaque glass tank (150 cm in length and 30 cm in breadth) filled with aquarium water of 10 cm depth. A 10-minute video recording of zebrafish swimming was made with a video camera (Sony, DSC-W830).

### 2.6. Histology and immunohistochemistry

Spinal cord tissues were dissected out and fixed in 4% paraformaldehyde (Sigma) for 8 hr or overnight at 4°C. Both injured and uninjured spinal cord tissues were embedded in O.C.T compound (Leica) and cryosectioned at 10 μm.

Immunofluorescence was performed in paraformaldehyde fixed 10 μm cryosections. Details of primary and secondary antibodies used for immunofluorescence were mentioned in the Key Resource Table. Briefly, sections were rehydrated and given several washes in Phosphate buffered saline (PBS) with 0.1% Tween-20 or Triton X 100 (PBST). The sections were first incubated with blocking solution for 1 hr (5% goat/rabbit/donkey serum, 1% BSA in PBST) and then with primary antibody for either 1 hr at room temperature or overnight at 4°C. Antigen retrieval was done wherever appropriate by keeping the slides in 80°C water bath for 15 min either in sodium-citrate buffer (pH 6.0) or Tris buffer (pH 8.0) before incubation with the primary antibody. This was followed by washes in PBST and incubated with secondary antibody for 2 hr at room temperature and nuclei were counter-stained with DAPI (Sigma).

After washing in PBS, the sections were mounted with Gelvatol mounting medium (Vector Labs, Burlingame, CA) and kept in the dark.

### 2.7. Axonal tracing

An incision has been made by fine scissor through the skin at approximately 3– 4 mm caudal to the spinal cord transection site. The spinal cord was exposed, and dry crystals of 1 ml of Fluoro-Ruby (tetra-methylrhodamine dextran, 10,000 MW; Thermo Fisher Scientific, USA) were placed on the surface of the exposed cord. The incision was closed, and the spinal cord was collected 24 hours after the dye insertion for histological analysis.

### 2.8. Quantitative reverse transcription polymerase chain reaction (qRTPCR) analysis

Total RNA from zebrafish spinal cord of different experimental groups was extracted using TRIzol reagent, and cDNA was subsequently synthesized using the GoScript™ Reverse Transcriptase Kit (Promega, Madison, USA). Primers were designed with web-based software Primer3 and primers were synthesized by Integrated DNA Technologies (IDT, Coralville, Iowa, USA). The primer sequences are mentioned in Table S1. qRT-PCR was performed using a LightCycler 480 real time machine (Roche, Switzerland). For RT-PCR, genes of interest were amplified using a PrimeSTAR GXL kit (Clontech, CA, USA). The amount of cDNA was normalized according to β-actin2 amplification in qRT-PCR and RT-PCR experiments. All qRT-PCR were performed using SYBR^TM^ Power Master Mix (Applied Biosystems, CA, USA) and results obtained from 3–5 biological replicates. The fold induction was calculated by setting control conditions to 1.

### 2.9. NAD assay

The cellular NAD^+^ content was quantified employing the cycling assay method, as outlined by Yang and Sauve (2021)(29). In brief, NAD^+^ extraction was carried out by homogenizing snap-frozen spinal cord tissues in 7% perchloric acid, followed by sonication of the mixture for 5 minutes on ice. The neutralization of NAD^+^ was achieved by adding 2N NaOH and 500 mM phosphate buffer. Subsequently, 1x cycling buffer was added to the wells of a 96-well microtiter plate. The cycling assay reagents, comprising 1x cycling buffer containing 54 μM, 2.25 mM L-lactate, and 0.4 U/mL Lactate Dehydrogenase, were then added to each sample well. Diaphorase was subsequently added to achieve a final concentration of 0.0035–0.007 U/mL in 1x cycling buffer, and the mixture was incubated for 10 minutes at room temperature. Optical density was measured with a kinetic fluorescence reading at 530 nm excitation and 580 nm emission, utilizing the Varioskan LUX multimode microplate reader (Thermo Fisher Scientific, Waltham, MA, USA).

### 2.10. ELISA

ELISA assay was done according to Dutta et al. (2009). Frozen spinal cord tissues from different experimental groups were rapidly homogenized in RIPA buffer and the tissue extracts were added to wells (65 mg protein/well) of the 96-well microtiter plate and incubated overnight at 4°C. After nonspecific sites were blocked with blocking buffer (1% BSA in PBS), the wells were incubated with primary antibody for 1 hr at 37°C, and washed thrice in washing buffer (0.5% BSA, 0.5% NP-40 in PBS). HRP coupled anti-goat antibody (1:1000 dilution) was used for incubating the plates at 37°C for 1 hr, then washed several times in washing buffer and substrate TMB (tetramethyl benzidine) was added in dark. The reaction was terminated by adding 1M H2SO4 as a Stop solution. The optical density was measured at 450 nm by using Varioskan LUX multimode microplate reader (Thermo Fisher scientific, Waltham, MA, USA).

### 2.11. Immunoprecipitation

Frozen spinal cord tissues from different experimental groups were rapidly homogenized in RIPA buffer and lysates were clarified by centrifugation for 20 minutes in a microcentrifuge at 4°C. After preclearing with 20 mL Protein A/G Sepharose beads for 1 hour at 4°C, tissue lysates were incubated with 5 µL primary antibody and rotated for 12 hours at 4°C. Protein A Sepharose beads (20 µL) were added to each sample and the samples were incubated for 2 hours at 4°C. The immune complex was washed three times with lysis buffer and was quantified using indirect ELISA method.

### 2.12. Quantification of Global DNA methylation

After isolating the genomic DNA from the spinal cord tissue using the Phenol-chloroform extraction method using Phenol:Choloroform:Isoamyl alcohol followed by ethanol precipitation, global DNA methylation was determined in the isolated genomic DNA using the Methyl Flash Methylated DNA Quantification Kit (Colorimetric) (Epigentek Group Inc., New York, NY, USA). The kit measures the methyl-cytosine content as a percentage of total cytosine content. The purified DNA in a concentration of 100 ng/μL was added to ELISA plate. The methylated fraction of DNA is quantified using 5-methylcytosine specific antibodies. The amount of methylated DNA was proportional to the OD intensity in an ELISA plate reader at 450 nm. DNA methylation was calculated using the formula: [(ODSAMPLE – ODNEGATIVE CONTROL)/S]/ [((ODPOSITIVE CONTROL – ODNEGATIVE CONTROL) x2)/P]x100; where OD is optical density; S is the amount of input sample DNA in ng; P is the amount of input positive control in ng. The amount of methylated DNA was expressed as percentage of total DNA.

### 2.13. Histone extraction from spinal cord tissue

Histone extracts were prepared by lysis of spinal cord tissues in PBS containing 0.5% Triton X-100, 2 mM phenyl-methylsulfonyl fluoride (PMSF), 0.02% NaN3, and incubated for 10 min on ice. The pellet was resuspended after centrifugation in 0.2 N HCl and incubated at 4°C overnight. After that samples were centrifuged and the supernatant was collected in a separate tube.

### 2.14. Quantification of Global H3 and H4 acetylation

Extracted histones were analyzed using the EpiQuick Total Histone H3 & H4 Acetylation Detection Fast Kit (Epigentek) according to the manufacturer’s protocols. Briefly, histone extract was added to the ELISA wells and incubated with primary antibody for 2 hours at room temperature. The wells were then washed and incubated with the secondary Ab-containing solution for 1 hour on an orbital shaker. The wells were then developed with the enzyme solution for 5 min and the absorbance was measured at 450 nm by using Varioskan LUX multimode microplate reader (Thermo Fisher scientific, Waltham, MA, USA).

### 2.15. Targeted Methylated DNA Immunoprecipitation (MeDIP) sequencing

Genomic DNA from spinal cords of different experimental conditions were isolated according to the previous method separately. Each condition had three biological replicates. Isolated genomic DNA was sonicated and three biological replicates of sonicated genomic DNA were pooled together for MeDIP-seq. Targeted sequencing was performed with Illumina HiSeq 2000 sequencing system. Low quality sequences were filtered off from raw sequencing data and high-quality pair-end MeDIP-seq sequences were aligned with the zebrafish genome (Zv10/danRer11) and methylated peaks were generated using WashU Epigenome browser (https://epigenomegateway.wustl.edu/browser/). Methylation percentage was determined from the ratio of heights of a methylated cytosine peak and the sum of methylated cytosine and unmethylated thymine peaks for each cytosine in a CpG dinucleotide (30). Quantification of CpG and non-CpG sites were performed using alignment reads.

### 2.16. Microscopy

Immunostained tissue sections were photographed by using an OLYMPUS BX53F2 microscope (Olympus, Tokyo, Japan). Confocal images were taken with a Zeiss LSM 710 confocal microscope (Carl Zeiss AG, Germany).

### 2.17. Quantification and Statistical analysis

#### 2.17.1. Cell Number Quantification

Cells positive with different markers (Sirt1^+^, Dnmt1^+^ and Yap1^+^) in regenerating spinal cord were imaged within the injured areas of the spinal cord (1360 x 1024 pixels) under 10x magnification and counted manually using ImageJ software (US National Institutes of Health, Bethesda, MD, USA). For each spinal cord, the number of cells was calculated by averaging the data from three particular sections.

#### 2.17.2. Swim Path Measurement

Total distance of swim path and the trajectories of the swimming movement were obtained from the video recording of swimming zebrafish using idTracker software (de Polavieja lab, Cajal Institute,Consejo Superior de Investigaciones Científicas, Madrid, Spain). The data were represented as mean ± SEM and Mann– Whitney U test was used for calculating P value.

#### 2.17.3. Precursor Cell Proliferation Quantification

To quantify NPC proliferation, transverse sections of the injury site, depicting the ependymal canal, were imaged (1360 x 1024 pixels) at 7 dpi, using an OLYMPUS BX53F2 microscope (Olympus, Tokyo, Japan). The numbers of cells positive for Sox2 and Sox2 co-expressing with PCNA were manually quantified utilizing ImageJ software. The proliferation index of neural progenitor cells in each spinal cord were calculated by averaging the percentages of Sox2^+^/ PCNA^+^ cells from three selected sections.

#### 2.17.4. Glial bridging Quantification

Glial bridging was calculated as a ratio of the cross-sectional area of the glial bridge (at the lesion site) and the area of the intact SC (300 µm rostral to the lesion) using ImageJ software.

#### 2.17.5. Statistical Analysis

All statistical values are represented as mean ± SEM. To determine the statistical significance, the P values were calculated either with Mann–Whitney U tests, Student’s t-tests, or Fisher’s exact tests and P values less than 0.05 considered as statistically significant. All qRT-PCR results were obtained from 3–5 biological replicates. Statistical methods, sample size and P values are also described in the respective figure legends.

## 3. Results

### 3.1. Injury-induced induction of *sirtuin1* among *sirtuin* family is associated with spinal cord regeneration

Our study aimed to explore the potential involvement of sirtuin-mediated epigenetic changes in the regeneration of the zebrafish spinal cord. To investigate the role of the sirtuin family of histone deacetylases in this process, we analyzed the mRNA expression of various members (*sirtuin1-7*) in the regenerating spinal cord of adult zebrafish at 7 dpi. Notably, we observed a significant upregulation of *sirtuin1* (*p*<0.01) and *sirtuin2* (*p*<0.05) during regeneration (Figure 1A and Supplementary Figure S1B). Importantly, *sirtuin1* (*sirt1*), the zebrafish ortholog of mammalian *Sirtuin1*, exhibited the most significant increase (*p<0.01*) compared to its expression in the uninjured spinal cord, as depicted in Figure 1A & S1B. This finding is consistent with the observation by Pereira et al. (2011), which identified *sirtuin1* as the most highly expressed member among all *sirtuin* family members in the adult zebrafish brain (31).

**Figure 1.**
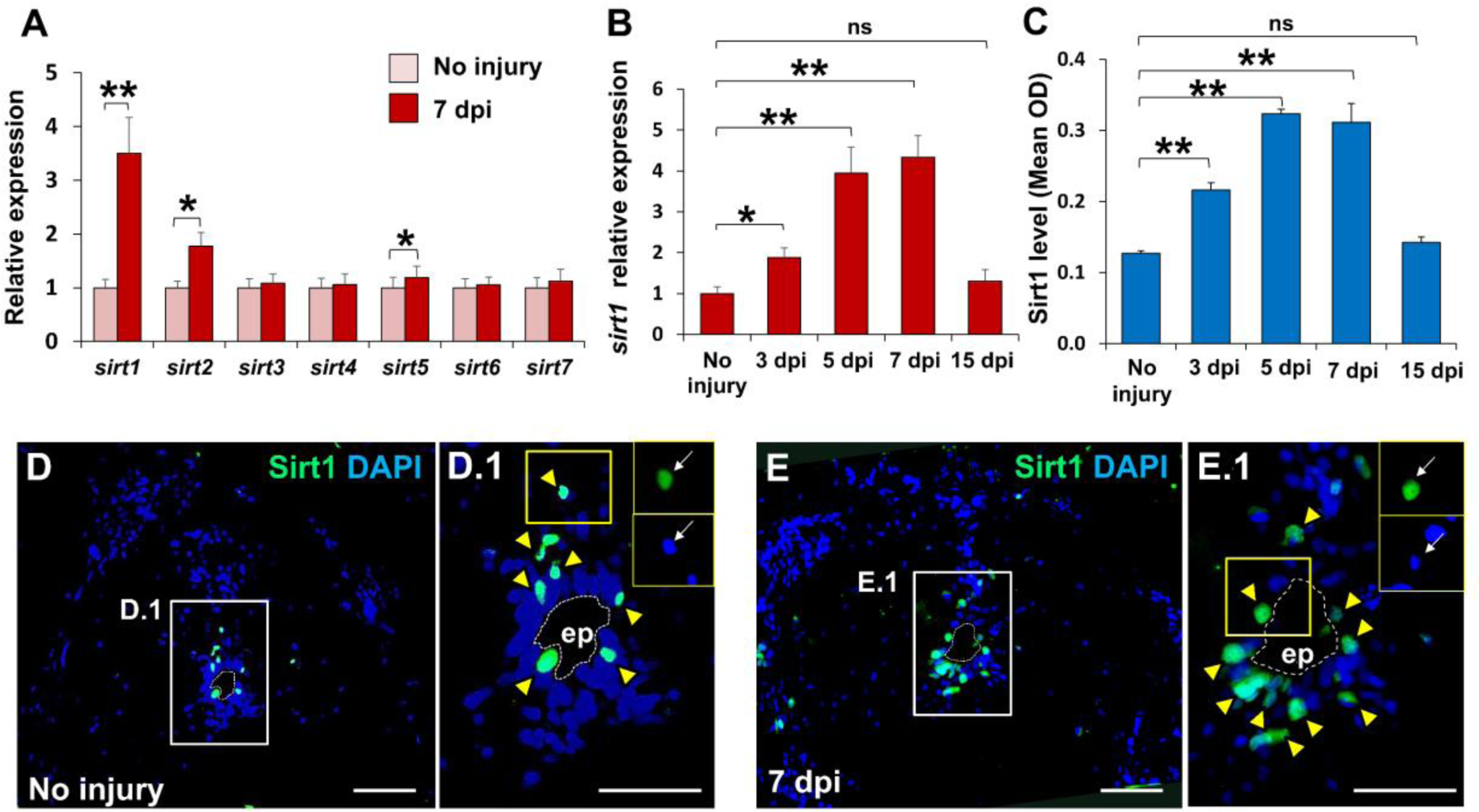
Legend: Injury-induced induction of *sirtuin1* expression among other sirtuin paralogs is associated with spinal cord regeneration in zebrafish. (A) qRT-PCR analysis of members of *sirtuin* family (Mean± SEM, n=5); (B) qRT-PCR analysis of *sirtuin1* mRNA time-course expression in regenerating spinal cord (Mean±SEM, n=5); (C) Quantitative analysis of Sirt1 protein level in different time points by indirect ELISA method (Mean±SEM, n=6); (D) Immunohistochemical staining of uninjured and 7 dpi injured spinal cord (transverse Section) showing expression of Sirt1. The white-dotted line marked the ependymal canal. Yellow arrowheads indicate the Sirt1^+^ cells. Magnification, 10x; Insets show single-channel slices of the demarcated region (magnification, 20x); Scale bar, 50µm.; ep, ependymal canal. \**p* < 0.05; \*\**p* < 0.01; ns, Non-significant; Mann-Whitney U test.

Following this observation, we examined the gene expression pattern of *sirt1* during the regeneration time course using both RT-PCR and semi-qRT-PCR methods. Both analyses revealed a significant upregulation of *sirt1* transcript as early as 3 dpi (*p<0.01*), reaching peak levels at 5 and 7 dpi (*p<0.01*), and subsequently declining by 15 dpi, coinciding with the nearly complete proliferation of neural progenitor cells (Figure 1B & S1C). In line with its mRNA expression pattern, the analysis of Sirt1 protein expression by indirect ELISA indicated a significant increase in Sirt1 activity around 5-7 dpi (*p<0.01*), followed by a reduction at later timepoints (non-significant) (Figure 1C). Immunohistochemical analysis demonstrated the presence of few Sirt1-positive cells in the uninjured spinal cord around the ependymal canal, with an increase in the number of Sirt1-positive cells after 7 dpi around the ependymal canal (Figure 1D and 1E).

To further investigate, we labelled neural progenitor cells after 7 dpi with the NPC marker Sox2 and observed that all Sox2-positive NPCs were also Sirt1-positive (Figure 2A and 2B). This observation suggests a close association between *sirt1* expression and injury-induced proliferating neural progenitor cells, as previously reported in white matter regeneration after neonatal brain injury in mice (32). In summary, our data indicate that *sirt1* is expressed in neural progenitor cells in the spinal cord after injury and is associated with zebrafish spinal cord regeneration.

**Figure 2.**
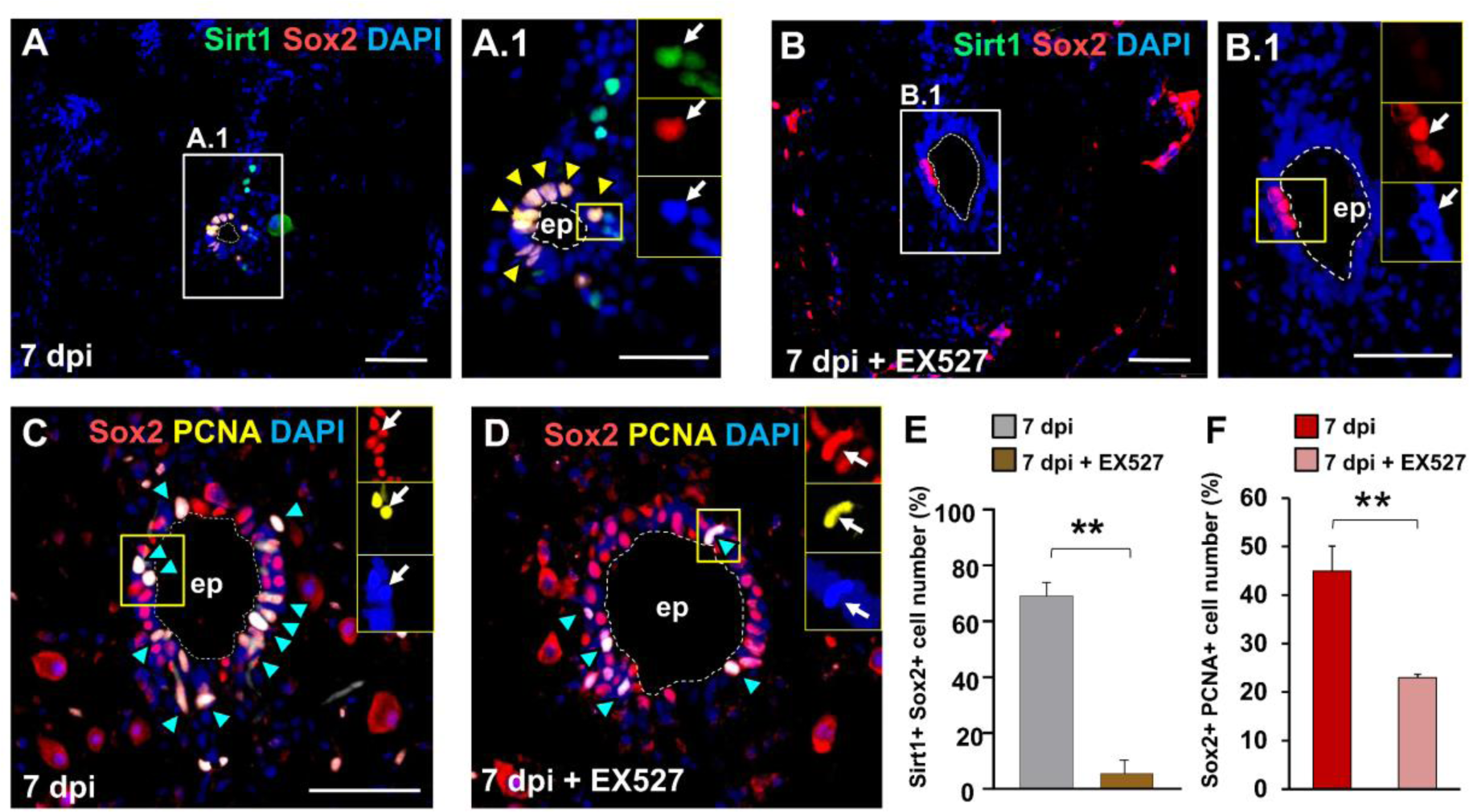
Legend: Sirt1 is required for proliferation of neural progenitor cells after SCI in zebrafish. (A, B) Immunohistochemistry of transverse section of zebrafish spinal cord showing co-expression of Sirt1 & Sox2 (marked by yellow arrowheads) after 7 dpi in both DMSO and EX527-treated condition. Magnification, 10x; Insets show single-channel slices of the demarcated region; Scale bar, 100µm. (C, D) Immunohistochemistry of transverse section of zebrafish spinal cord showing Proliferating neural progenitor cells (co-expression of Sox2 & PCNA, marked by blue arrowheads) after 7 dpi in both DMSO and EX527-treated condition. The white-dotted line marked the ependymal canal. Magnification, 20x; Insets show single-channel slices of the demarcated region. Scale bar, 50 µm. (E) Quantification of (A, B); (F) Quantification of (C, D). Data represented as Mean± SEM, n=5; \*\**p* < 0.01; Mann-Whitney U test.

### 3.2. Sirt1 induces proliferation of neural progenitor cells in injured spinal cord

To determine whether Sirt1 plays a specific role in NPC proliferation in regenerating spinal cord, we analyzed spinal cord lesions at 7days after transection, that is, at a time point when NPC proliferation is maximal. To investigate whether Sirt1 is required for NPC proliferation in adult zebrafish spinal cord after injury, we functionally inhibited the Sirt1 activity by using Sirt1-selective inhibitor, EX527. The inhibitor, EX527 was applied by local intraspinal injection at the injury site. Indirect ELISA analysis showed that EX527-mediated functional inhibition resulted in significant decrease (*p<0.01*) in Sirt1 protein level in regenerating spinal cord (Figure S1D). To observe whether Sirt1 was selectively and functionally inhibited by EX527, NAD^+^ quantification was assessed after inhibitor treatment by cyclic NAD assay. As NAD^+^ is the cofactor required for Sirt1 activity, analysis of intracellular NAD^+^ level during spinal cord regeneration showed a significantly increased level throughout the regeneration time course while in case of EX527 application, intracellular level of NAD^+^ significantly decreased (*p*<0.01, Figure S1E). As EX527 requires NAD^+^ for inhibition, therefore from the intracellular level of both Sirt1 and NAD^+^ in case of EX527-treatment it can be confirmed that EX527 selectively inhibit the Sirt1 activity. As previously observed from immunohistochemical study, the number of both Sirt1-positive and Sox2-positive cells increased after 7 dpi around the ependymal canal. After EX527 treatment, there were significant decrease (*p<0.01*) in the number of both Sirt1-positive and Sox2-positive cells around the ependymal canal at around 7 dpi, indicating that EX527 reduces the Sirt1 expression after injury.

Next, we assessed the effect of Sirt1 inhibition on NPC proliferation in regenerating spinal cord after 7 dpi by immunolabeling with the neural progenitor marker, Sox2 and the proliferation marker PCNA. In EX527-treated regenerating spinal cord, after 7 dpi, a significant reduction (*p<0.01*) in both Sox2-positive and PCNA-positive proliferating neural progenitor cells was observed. We also observed a significant (*p<0.05*) decrease in the percentage of Sox2^+^/PCNA^+^ NPCs by translational blocking of endogenous *sirt1* mRNA by using *sirt1* antisense morpholino (Figure S2b & S2C). These data indicate that EX527-mediated Sirt1 inhibition displayed a ∼37% reduction in Sirt1 expression as well as a ∼13% reduction in NPC proliferation which is extremely significant (*p<0.01*), thereby suggesting a positive role of Sirt1 in NPC proliferation after injury.

### 3.3. Inhibition of Sirtuin1 impedes functional recovery after spinal cord injury

Accumulating evidence highlights the role of Sirt1 in regulating various neurological functions in mammals through the deacetylation of many proteins besides histones. Studies in mice demonstrated that genetic inhibition of Sirt1 impeded axonal development in embryonic hippocampal neurons, while both genetic and pharmacological upregulation of Sirt1 not only facilitated axon formation but also promoted axon elongation (33). SIRT1 has been recognized for its central role in axonogenesis and optic nerve regeneration, as evidenced by previous studies (34–36). However, the existing evidences regarding the interplay between Sirt1 and axonal regrowth in zebrafish remains limited.

NPC proliferation constitutes a crucial prerequisite for the formation of glial bridges and axonal regeneration following spinal cord injury, therefore we conducted experiments to assess the impact of Sirt1 in spinal cord regeneration, focusing specifically at glial bridging, axon regrowth, and swim behaviour. For glial bridging, GFAP immunostaining was employed to quantify the areas of glial bridges at the lesion site relative to intact spinal cord tissues. Here, we observed robust glial bridging in DMSO-treated control animals at 21dpi (Figure 3A). Conversely, animals treated with EX527 displayed a substantial ∼71% reduction (*p<0.01*) in bridging compared to the control group (Figure 3A and 3D). This reduction may be attributed to the observation that, at 21 dpi, glial cells within the injury sites of Sirt1-inhibited zebrafish often failed to extend projections into the lesion. As it has been observed earlier that EX527-treated zebrafish displayed a ∼13% reduction in NPC proliferation at 7 dpi (Figure 2F) it suggests that the effects of Sirt1 inhibition were preferential to progenitor cells and therefore, Sirt1 is required for the changes in NPC proliferation and morphology of glial cells during spinal cord regeneration.

**Figure 3.**
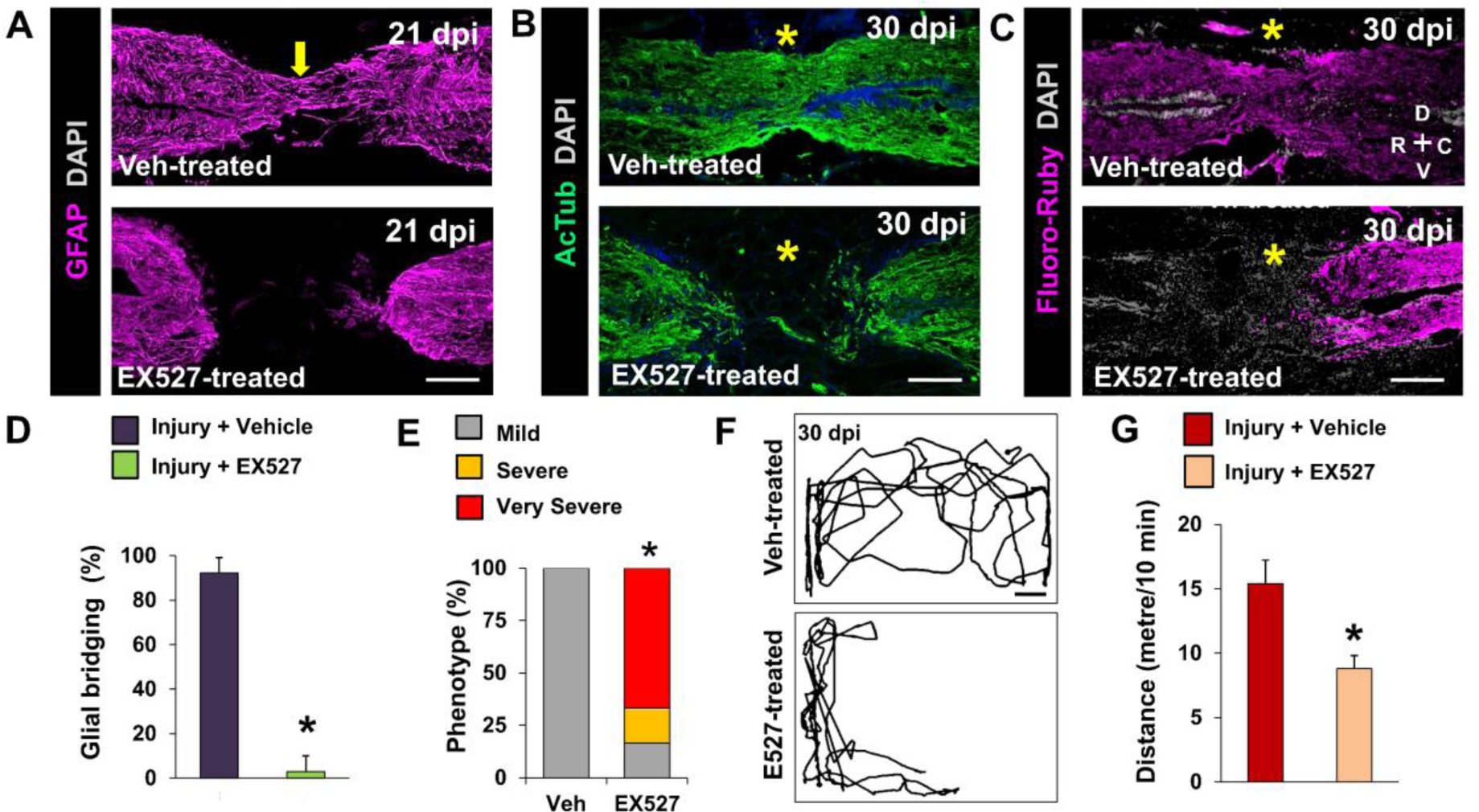
Legend: Sirt1 is required for axonogenesis after SCI in zebrafish. (A) Immunohistochemical staining of spinal cord sections after 21 dpi from DMSO-treated (Upper) and EX527-treated (Lower) fish showing glial bridge formation; (B) Immunohistochemical staining of spinal cord sections after 30 dpi from DMSO-treated (Upper) and EX527-treated fish showing axonal projections; (C) Retrograde tracing of axonal projections using Fluoro-Ruby (FR). Regenerating axons spanned the transection injury in DMSO-treated fish (Upper), but not in EX527-treated fish (Lower); (D) Quantification of glial bridging in (A) (N=5); (E) Quantification of regeneration in (B) (N=5); (F) Swim tracking of individual animals after 30 dpi from DMSO-treated (Upper) and EX527-treated (Lower) experimental groups (N=8); (G) Quantification of total distance moved in (F). Yellow arrow indicates glial bridging; Asterisks indicate injury epicentre; Confocal projections of z stacks are shown. \**p* < 0.05; \*\**p* < 0.01; Fisher’s exact test (E) and Mann-Whitney U test (D, G). Scale bars, 100 µm (A, B, C) and 0.1 m (F).

Utilizing the functional inhibition of Sirt1 by applying EX-527, we assessed axonal regrowth by immunostaining axonal fibres with an anti-Acetylated tubulin antibody following spinal cord transection in both DMSO-treated and EX527-treated zebrafish (Figure 3B). In the case of DMSO-treated fish, the majority of injured spinal cords regenerated, with no visible gap at the injury site by 30 dpi (Figure 3B, Mild in 3E). Conversely, at 30 dpi, EX527-treated zebrafish exhibited rostral and caudal stumps with disorganized axonal sprouts (Figure 3B, Severe in 3E), and around two-thirds of the Sirt1-inhibited zebrafish displayed disconnected rostral and caudal stumps at 30 dpi (Figure3B, very severe in 3E), as confirmed by retrograde tracing of axonal projections (Figure 3C).

While vehicle-treated control fish had recovered their pre-injury swimming ability by 30 dpi, EX527 treatment significantly impeded functional recovery, as evidenced by locomotor activity measurements (Figures 3F and 3G). Fish from the Sirt1-inhibited group exhibited approximately a 43% reduction in swimming ability (*p<0.05*), as measured by the travelled swimming distance, compared to the DMSO-treated control group. In case of *sirt1* antisense morpholino treatment, we observed similar kind of incomplete glial bridging (Figure S2D) as well as significantly reduced swimming activity (Figure S2E & S2F) which resulted in poor functional recovery in comparison to control group. Previous observations suggested that fish with high glial bridging tended to swim longer distances, and swim distance assessed during the early phase of spinal cord regeneration could predict injury outcomes at 8 weeks post-injury (wpi) (37). Consistent with this, our study noted that fish from the DMSO-treated control group, displaying significant glial bridging, travelled much longer distances. Simultaneously, the control group exhibited substantial axonal regeneration, whereas EX527-treated fish displayed incomplete glial bridging, covered less distance, and subsequently exhibited significantly reduced axonal regeneration. Taken together, these data demonstrated that inhibition of Sirt1 caused a regenerative defect in the zebrafish spinal cord and thus Sirt1 promotes functional recovery after SCI by controlling the changes in proliferation and morphology of glial cells and subsequent axonal regeneration.

### 3.4. Resveratrol treatment induces NPC proliferation and axonal regrowth in zebrafish spinal cord by activating Sirt1 expression

Prior studies have suggested that resveratrol does not alter mRNA level of the Sirt1 gene in zebrafish liver following exposure (38). Another study reported an opposite observation that resveratrol increased Sirt1 gene and protein expression in the zebrafish retina (39). In our study, we observed a significant increase (*p<0.01*) in Sirt1 protein level at 3, 5,7 and 15 dpi along with uninjured fish when compared to the EX527-treated Sirt1-inhibited fish (Figure 4A). Next, we observed the rescue effect of resveratrol on the Sirt1-mediated induction in the NPC proliferation after injury and immunohistochemical analysis using Sox2 and PCNA markers revealed an increased percentage of proliferating neural progenitor cells upon Sirt1-induction by resveratrol treatment (Figure 4B & 4C). We also found that resveratrol treatment induced the glial bridge formation when compared to the EX527-mediated Sirt1-inhibited condition (Figure 4D). We assessed axonal regrowth by immunostaining axonal fibres with an anti-Acetylated tubulin antibody (Figure 4E) and found that majority (almost 83%) of the resveratrol-treated zebrafish showed regenerated axons with no visual gaps between the rostral and caudal stumps (Figure 4E, Mild in 4F) in comparison to the EX527-treated fish where almost two-third of the fish showed disconnected rostral and caudal stumps (Figure 4E, very severe in 4F). All these observations suggest that resveratrol treatment successfully rescued the inhibitory effect of EX527 on Sirt1 activity.

**Figure 4.**
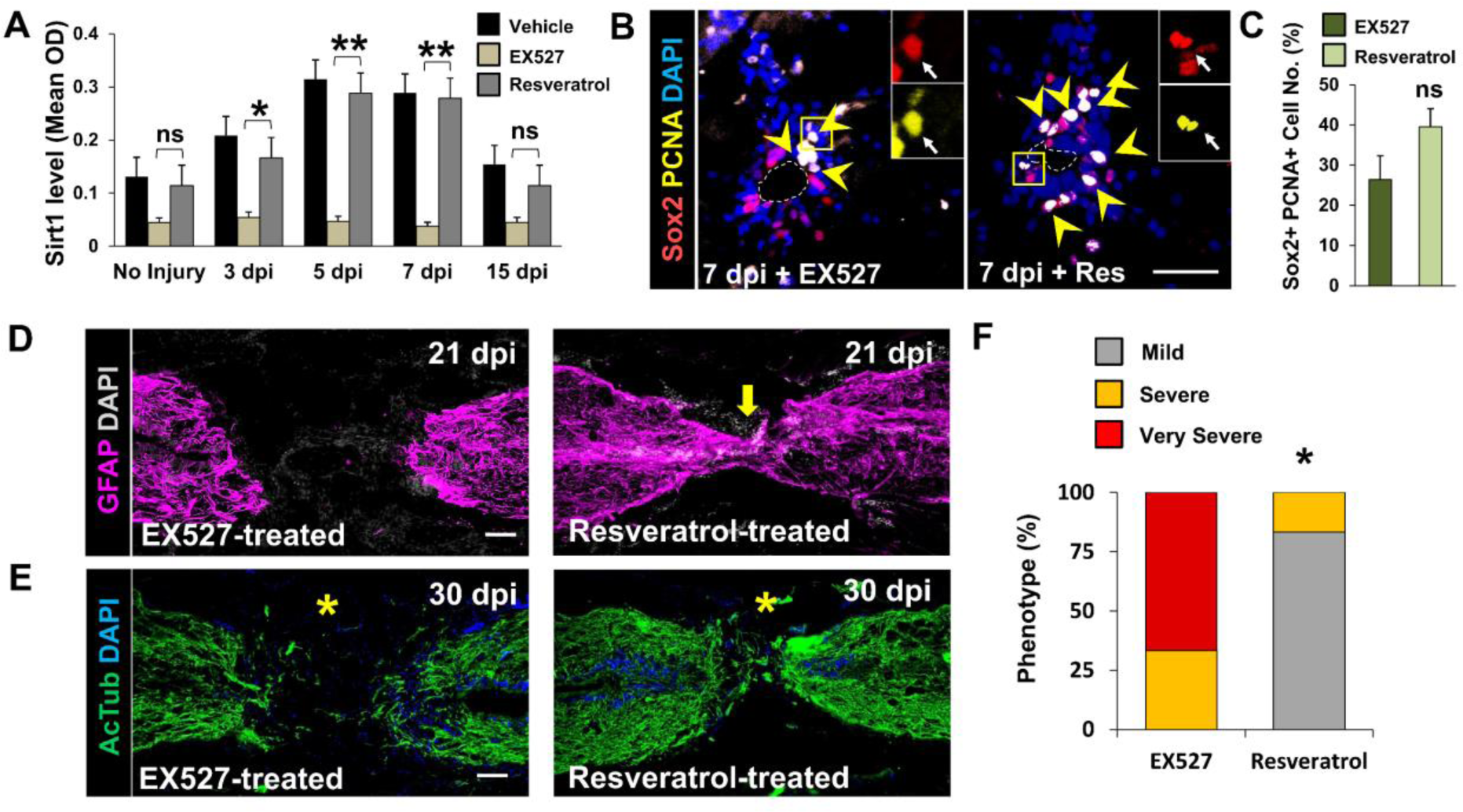
Legend: Resveratrol induces NPC proliferation and axonal regrowth by activating Sirt1 expression after SCI in zebrafish. (A) Quantitative analysis of Sirt1 protein level in different time points by indirect ELISA method from DMSO-treated, EX527-treated and Resveratrol-treated fish, (Mean±SEM, n=6); (B) Immunohistochemistry of transverse section of zebrafish spinal cord showing Proliferating neural progenitor cells (co-expression of Sox2 & PCNA, marked by yellow arrowheads) after 7 dpi in both EX527-treated (left) and Resveratrol-treated (right) condition. The white-dotted line marked the ependymal canal. Magnification, 20x; Insets show single-channel slices of the demarcated region; (C) Quantification of (B), (N=5); (D) Immunohistochemical staining of spinal cord sections after 21 dpi from EX527-treated (left) and Resveratrol-treated (right) fish showing glial bridge formation; (E) Immunohistochemical staining of spinal cord sections after 30 dpi from EX527-treated (left) and Resveratrol-treated (right) fish showing axonal projections; (F) Quantification of regeneration in (E). Yellow arrow indicates glial bridging; asterisk indicates injury epicentre. Confocal projections of z stacks are shown. Data represented as Mean±SEM, n=5; \**p* < 0.05; ns, non-significant; Mann-Whitney U test (C) and Fisher’s exact test (F). Scale bar, 50 µm.

### 3.5. Sirt1 regulated global methylation level during spinal cord regeneration

The accessibility of regeneration-associated genes (RAGs) for transcription remains high and increases with injury in regenerating species (40,41). Alteration in DNA methylation is one of the mechanisms associated with the regulation of the accessibility to RAGs. In Xenopus, elongation factor 1 *α* (*ef1α*) became demethylated and subsequently activated after caudal fin amputation or retinal injury while during homeostatic condition it became methylated and remained inactive (12). To observe the change in the global DNA methylation during zebrafish spinal cord regeneration, we quantified the percentage of 5-methylated cytosine (5-mC%) in zebrafish spinal cord during regeneration time course (3, 7 and 15 dpi). It has been observed that 5-mC% displayed a 1.76% reduction at 3dpi and 3.13% reduction at 7 dpi when compared to uninjured spinal cord but also started to increase significantly at 15 dpi exhibiting 2.72% significant increment (*p<0.01*, Figure 5A). Sirt1 inhibition demonstrated higher DNA methylation peak at 7 dpi (∼1% increase) (*p<0.01*), but no significant changes in the 5-mC% at 3 and 15 dpi (Figure 5A), thereby giving a better insight about the modulating role of Sirt1 in the methylation status at the DNA level when NPC proliferation reaches its peak during regeneration.

**Figure 5.**
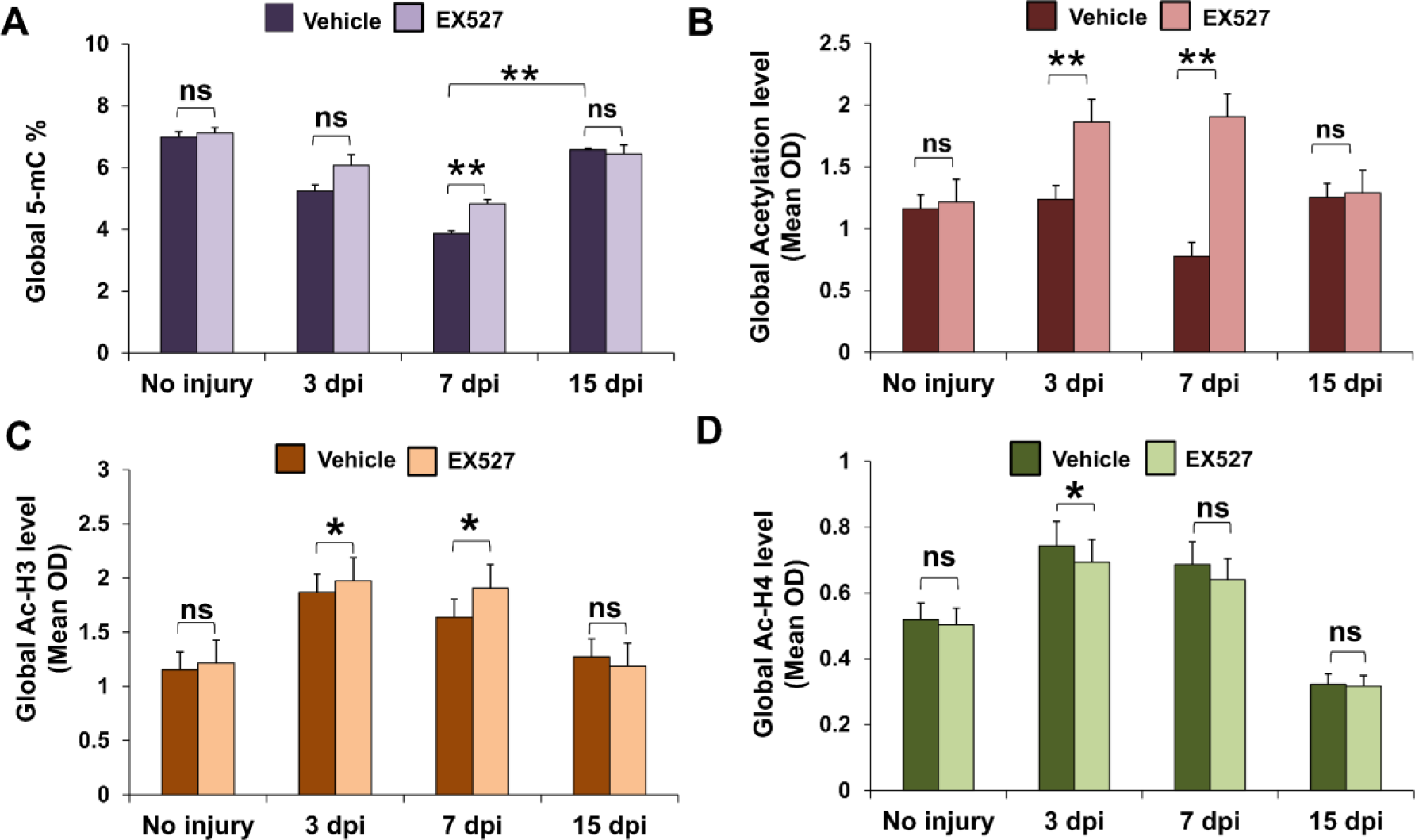
Legend: Sirt1 mediates global DNA methylation and acetylation level during spinal cord regeneration in zebrafish. (A) Quantitative analysis of global 5-mC percentage in different time points by indirect ELISA method from DMSO-treated and EX527-treated fish; (B) Quantitative analysis of global acetylation level in different time points by indirect ELISA method from DMSO-treated and EX527-treated fish; (C) Quantitative analysis of global Acetylated-H3 level in different time points by indirect ELISA method from DMSO-treated and EX527-treated fish; (D) Quantitative analysis of global Acetylated-H4 level in different time points by indirect ELISA method from DMSO-treated and EX527-treated fish. Data represented as Mean±SEM, n=6; \**p* < 0.05; \*\**p* < 0.01; ns, Non-significant; Mann-Whitney U test.

### 3.6. Sirt1 modulates the global deacetylation of H3 histone proteins but not H4 during spinal cord regeneration

Zebrafish Sirt1 is a member of NAD^+^-dependent protein deacetylases, with a vast list of non-histone proteins including the histone protein H3 and H4 (42–44). To determine their involvement in Sirt1-induced deacetylation in global landscape, we investigated the effect of functional inhibition of Sirt1 on the global acetylation levels in histone extract during regeneration time course using anti-pan acetylation antibody (Figure 5B). It reflects that the global acetylation level has been decreased gradually during the regeneration time course showing its lowest level at 7 dpi while in case of Sirt1-inhibited condition the global acetylation levels at 3 dpi and 7 dpi have been increased significantly (*p<0.01*, Figure 5B). This observation supports the Sirt1-directed deacetylation during early time-points of regeneration (3 dpi and 7 dpi) which was absent in case of Sirt1-inhibited condition inducing the global acetylation level up to 1-fold.

Next, we investigated the role of Sirt1 in global level of H3 and H4 acetylation using control (treated with DMSO) and Sirt1-inhibited zebrafish spinal cord during specific time points of regeneration event (Figure 5C and 5D). In case of global histone acetylation, when compared to the uninjured group, the global histone H3 acetylation level throughout the regeneration time course was significantly higher with the highest peak at around 3 and 7 dpi (*p<0.05*). While the EX527 treatment further increased the total histone H3 acetylation levels, our data did not show any significant differences in acetylation of histone H4 at the time studied among vehicle treated and EX527-treated animals except at 3 dpi (*p*<0.05).

### 3.7. Sirt1-mediated deacetylation regulates *dnmt1* expression during regeneration

DNA methylation plays a pivotal role as a regulatory mechanism during cell dedifferentiation and proliferation in the context of regeneration. While global alterations in DNA methylation have been documented during fin and retina regeneration(45,46), this phenomenon has not yet been explored in the context of spinal cord regeneration. Within the DNA methylation regulatory machinery, DNA methyltransferases (Dnmt) assume a central role, and among the various members of the Dnmt family, Dnmt1 stands out as a key modulator responsible for maintaining methylation patterns and facilitating gene silencing(47). Beyond its role in DNA methylation, DNMT1 also exerts transcriptional repression through methylation-independent mechanisms. For instance, it recruits transcription corepressor DNMT1-associated protein 1 (DMAP1), HDAC1, HDAC2, and methyl-CpG-binding protein to DNA (48,49).

In this study, we noted a transient reduction in the level of 5mC% at 7 dpi, followed by a gradual recovery during late phases of spinal cord regeneration (Figure 5A). This observation led us to infer the involvement of DNA demethylation processes in SC regeneration. To investigate which *dnmt* paralogs in zebrafish are involved in DNA demethylation during SC regeneration, we observed the mRNA expression patterns of all *dnmt* paralogs in zebrafish (*dnmt1*, *dnmt2*, *dnmt3aa*, *dnmt3ab*, *dnmt3ba*, *dnmt3bb.1*, *dnmt3bb.2* and *dnmt3bb.3*) in regenerating spinal cord (Figure 6A) by RT-PCR. Among the examined members of *dnmt* family, only two, namely *dnmt1* and *dnmt2*, displayed significant changes in the mRNA expression in the 7 dpi regenerating spinal cord. Though *dnmt2* (also known as *tRNA aspartic acid methyltransferase 1*) is primarily responsible for tRNA^ASP^ methylation (50), we emphasized our investigation on the other *dnmt* family member, *dnmt1*, which exhibited distinct and extremely significant change (*p<0.01*) in mRNA expression in the regenerating cord at 7 dpi (Figure 6A). This observation suggests that the specific paralog, *dnmt1,* may play a role in regulating 5mC levels and spatio-temporal gene expression during SC regeneration. Previous studies have indicated that inhibiting DNMT1 could result in reduced methylation levels in various animals, including frogs, mice, and humans(51–54). This implies a critical role for DNMT1 in maintaining global DNA methylation levels. In our investigation, we can indicate a correlation between the reduced expression of *dnmt1* and the decrease in global 5-mC% at 7 dpi. This suggests that, among the predominantly expressed *dnmt* paralogs in zebrafish, *dnmt1* is more positively associated with regulating the global methylation pattern during regeneration.

**Figure 6.**
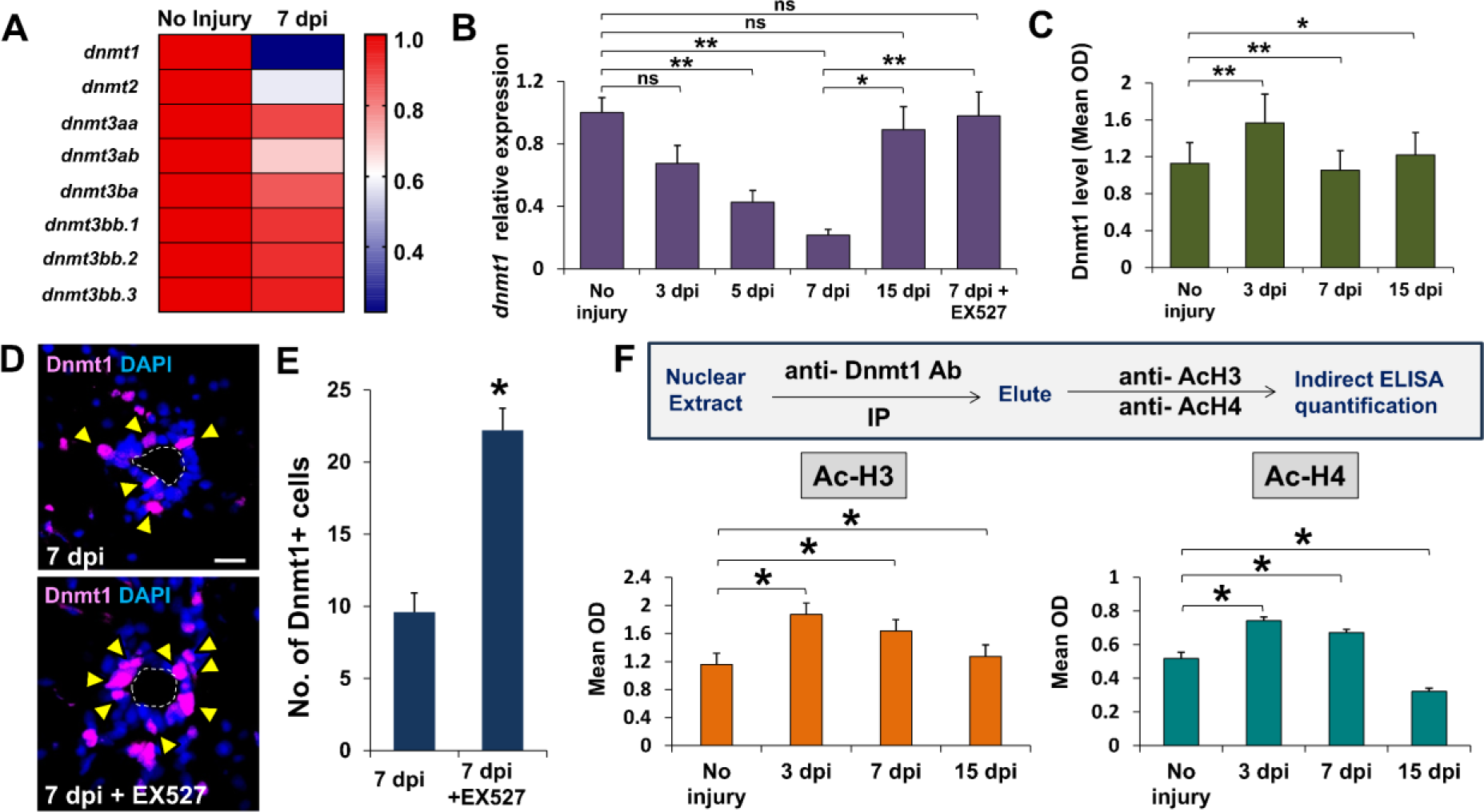
Legend: sirt1-mediated deacetylation regulates dnmt1 expression during spinal cord regeneration. (A) Heatmap visualization of mRNA expression of members of *dnmt* family by qRT-PCR analysis (Mean±SEM, n=5). A scale is shown on the right, in which red and blue correspond to a higher and a lower expression, respectively; (B) qRT-PCR analysis of *dnmt1* mRNA time-course expression in regenerating spinal cord (Mean±SEM, n=5); (C) Quantitative analysis of Dnmt1 protein level in different time points by indirect ELISA method (Mean±SEM, n=6); (D) Immunohistochemical staining of 7 dpi injured spinal cord (Transverse Section) in DMSO-treated (Upper) and EX527-treated (Lower) experimental groups showing expression of Dnmt1. The white-dotted line marked the ependymal canal, yellow arrowheads indicate Dnmt1^+^ cells. Magnification, 20x; Insets show single-channel slices of the demarcated region; Scale bar, 50µm; (E) Quantification of (D). (F) Quantitative analysis of Dnmt1 specific acetylated-H3 and acetylated-H4 level in nuclear extract from spinal cord tissues by immunoprecipitation followed by indirect ELISA method. Data represented as Mean±SEM, \**p* < 0.05; \*\**p* < 0.01; ns, Non-significant; Mann-Whitney U test.

To further investigate the temporal expression dynamics of *dnmt1* during the regeneration process, we examined the mRNA expression pattern of *dnmt1* at distinct time-points, specifically 3, 5, 7, and 15 dpi (Figure 6B and S3A). Both RT-PCR and semi-qRT-PCR methods were employed for this analysis, and both approaches consistently demonstrated a gradual reduction in the expression of *dnmt1* transcript up to 7 dpi, where the expression became nearly absent. Notably, at the 15 dpi time-point, the expression exhibited a substantial increase, surpassing levels observed in the uninjured control spinal cord (Figure 6B & S3A). Concurrently, the assessment of Dnmt1 protein expression through the indirect ELISA method revealed a parallel trend. The Dnmt1 activity demonstrated a gradual and significant decrease up to 7 dpi (*p<0.01*), followed by a notable increase during the later time-point at 15 dpi (*p<0.05*) (Figure 6C). These findings collectively underscore the dynamic and intricate regulatory pattern of *dnmt1* expression and activity throughout the spinal cord regeneration time course.

As we noted a significant elevation in the global 5-mC% levels at 7 dpi following Sirt1 inhibition, this may indicate a potential regulatory role of Sirt1 in governing DNA methylation during spinal cord regeneration in a global landscape. Intriguingly, EX-527-mediated Sirt1 inhibition resulted in an enhancement of *dnmt1* transcript levels specifically at 7 dpi (Figure 6B & S4A).

To validate the Sirt1-directed regulation, we subsequently quantified endogenous Dnmt1 level using the indirect ELISA method on EX527-treated regenerating spinal cord tissues (Figure S4B). Impressively, relative to the untreated control, Dnmt1 level in fish treated with EX-527 exhibited markedly elevated activities throughout the regeneration time-course, specifically at 3 dpi (*p<0.05*),7 dpi (*p<0.01*) and 15 dpi (*p<0.05*). Immunohistochemical analysis further corroborated these findings, indicating a limited presence of Dnmt1-positive cells in the regenerating spinal cord at 7 dpi around the ependymal canal (Figure 6D). Conversely, the number of dnmt1-positive cells significantly increased (*p<0.05*) under Sirt1-inhibited conditions at 7 dpi (Figure 6D and 6E). These outcomes collectively underscore the pivotal role of Sirt1 in orchestrating the regulation of both dnmt1 expression and Dnmt1 activity during spinal cord regeneration.

As further evidence of a stimulatory effect of deacetylation on Dnmt1 activity and as evidence that the effect is mediated by Sirt1, we conducted immunoprecipitation from nuclear extracts using an anti-Dnmt1 antibody. Subsequently, acetylated-H3 (ac-H3) and acetylated-H4 (ac-H4) levels were quantified in the elute utilizing the indirect ELISA method. In the detection of acetylated-H3 levels, a comparison throughout the regeneration time course revealed a relatively higher ac-H3 level at the early time point, diminishing at 7 dpi (Figure 6F). Under Sirt1-inhibited conditions, the ac-H3 level exhibited a significant increase during the regeneration time-course, peaking predominantly at 7 dpi (*p*<0.01) (Figure S4C). However, acetylated-H4 levels remained unaffected by Sirt1 inhibition throughout the regeneration time-course when compared to the DMSO-treated control condition (Figure 6F & S4D). This observation suggests that the previously noted reduction in Dnmt1 activity due to Sirt1 inhibition is likely a consequence of Sirt1-directed deacetylation of Dnmt1, with deacetylation being primarily associated with the H3 protein rather than H4. Taken together, these findings indicate that Sirt1 deacetylates the H3 histone protein in Dnmt1 during regeneration. By examining both Sirt1 expression and the acetylation level of Dnmt1 at 7 dpi, a time point coinciding with the induction of NPC proliferation, we can hypothesize that this Sirt1-mediated deacetylation and subsequent inactivation of Dnmt1 play a crucial role in controlling NPC proliferation.

### 3.8. Sirt1-mediated induced activation of Yap1 through hypomethylation and its nuclear translocation

As documented by various researchers (7,55), and consistent with our own findings, NPCs undergo epithelial-to-mesenchymal transition (EMT) following injury. During the initial stages, the expression of epithelial markers (i.e. E-cadherin), remains elevated. However, as the regenerative process unfolds, there is a notable shift towards increased expression of mesenchymal genes, including *vimentin* and *twist1*, concomitant with a decline in epithelial marker expression. Mechanistically, this EMT phenomenon is orchestrated at the transcriptional level through processes involving DNA methylation, histone modifications, and RNA-mediated epigenetic regulation (56–58). Given our emphasis on Sirt1 as an epigenetic regulator in the context of NPC proliferation, we observed the role of Sirt1 during EMT by functionally inhibiting Sirt1 in the zebrafish spinal cord post-injury, with a focus on the expression of *vimentin* as a representative mesenchymal marker. Notably, upon Sirt1 inhibition, there was a discernible reduction in Vimentin expression, particularly in proximity to the injury epicentre, contrasting with DMSO-treated control fish where Vimentin expression was more pronounced near the injury epicentre (Figure S5). This observation suggests a potential affirmative regulatory role of Sirt1 in the EMT process.

As Glial Epithelial-to-Mesenchymal Transition (EMT) is intricately connected to Yap1-Ctgfa signalling or HIPPO signalling pathway (59)(60) and considering our previous findings indicating a potential positive regulatory role of Sirt1 in EMT during regeneration, we propose that the activation of Sirt1 during regeneration leads to the activation of Yap1, subsequently resulting in the upregulated expression of *ctgfa*. This cascade of events is anticipated to induce ERG cell proliferation and, ultimately inducing glial bridging.

To investigate the influence of Sirt1 on the activation of Yap1, an indirect ELISA was conducted utilizing a Yap1-specific antibody. This assay aimed to assess Yap1 level in tissue lysates obtained from both uninjured and injured spinal cords at various regeneration time points (3, 5, 7, and 15 dpi) under two experimental conditions: DMSO-treated control and Sirt1 inhibition mediated by EX527. In the DMSO-treated control condition, a progressive increase in Yap1 level was observed, reaching significantly higher peaks at 5 and 7 dpi (*p<0.05*). Conversely, under Sirt1-inhibited condition, Yap1 activity exhibited a significant decrease (*p<0.01*) across all regeneration time points (Figure 7A).

**Figure 7.**
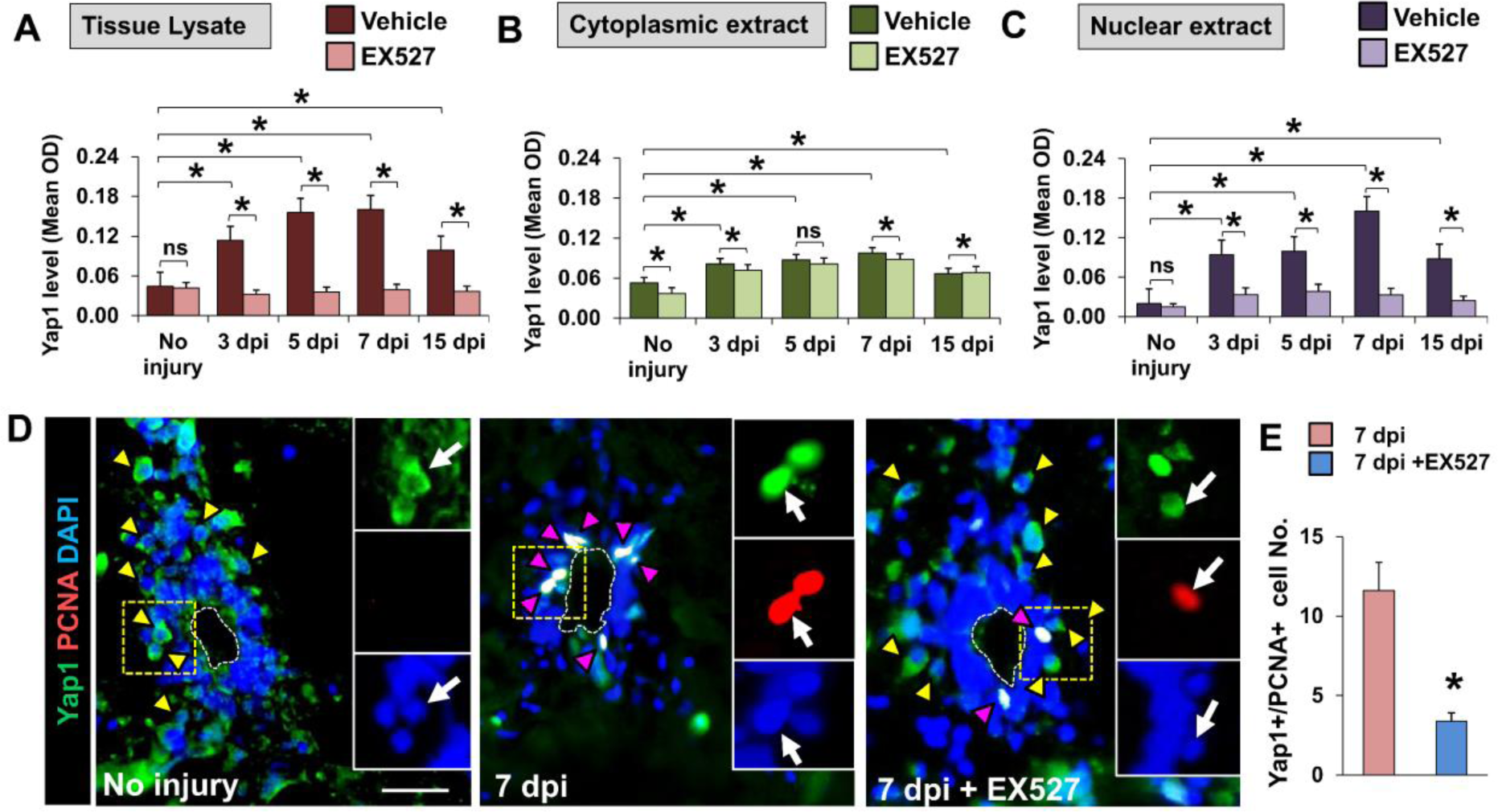
Legend: sirt1-mediated activation of Yap1 during spinal cord regeneration. (A, B, C) Time-course specific quantitative analysis of Yap1 protein level in tissue lysate (A), cytoplasmic extract (B) and nuclear extract (C) by indirect ELISA method in DMSO-treated and EX527-treated experimental groups (Mean±SEM, n=5). (D) Immunohistochemical staining of 7 dpi injured spinal cord (Transverse Section) in uninjured (left), DMSO-treated (middle) and EX527-treated (right) experimental groups showing Yap1 expression (marked by yellow arrow-heads) and co-expression of Yap1 and PCNA (marked by magenta arrow-heads). The white-dotted line marked the ependymal canal. Magnification, 20x; Insets show single-channel slices of the demarcated region; White arrow indicates a particular cell in the demarcated area in each single-channel slices; Scale bar, 50 µm. (E) Quantification of Yap1^+^ /PCNA^+^ cell numbers in (D) Data represented as Mean±SEM, *p < 0.05; **p < 0.01; ns, Non-significant; Mann-Whitney U test.

In the context of regeneration following injury, the HIPPO pathway undergoes inactivation, leading to the activation of Yap1. Subsequently, Yap1 undergoes cytoplasm-to-nuclear translocation, initiating the transcription of pro-proliferative genes, such as *ctgfa* (59). Henceforth, we postulate that Sirt1 facilitates Dnmt1-mediated demethylation or hypomethylation of Yap1 at the transcriptional level, thereby activating Yap1 and promoting its nuclear translocation. In order to substantiate this hypothesis, an additional experiment was conducted to investigate whether Sirt1 positively influences Yap1 activation and its nuclear translocation where Yap1 activity was assessed using an anti-Yap1 antibody in both cytoplasmic and nuclear extracts obtained from the same tissue lysate under various experimental conditions (Figure 7B and 7C). In the uninjured spinal cord, Yap1 activity in the cytoplasmic extract surpassed that in the nuclear extract, suggesting cytoplasmic retention of Yap1 in conditions characterized by low Sirt1 expression. Within the cytoplasmic extract, Yap1 activity exhibited a gradual increase during the early regeneration time course (3, 5, and 7 dpi), peaking significantly at 7 dpi (*p<0.05*). Conversely, under Sirt1 inhibition, no very significant reduction was observed at 3, 5, and 15 dpi, but a significant decrease in Yap1 activity occurred at 7 dpi when compared to untreated control fish (*p*<0.05).

A similar trend was observed in DMSO-treated injured fish during the regeneration time course, with Yap1 activity in the nuclear extract gradually increasing, reaching a higher peak at 7 dpi (*p<0.05*), and subsequently decreasing at 15 dpi (*p<0.05*) (Figure 7C). However, Yap1 activity significantly decreased in Sirt1-inhibited conditions during the regeneration time course in the nuclear extract, indicating cytoplasmic retention instead of nuclear translocation during the low Sirt1 activity.

Immunohistochemical analysis, employing antibodies against Yap1 and PCNA, provided further support for this finding (Figure 7D). In the uninjured spinal cord, cells with cytoplasmic expression of Yap1 were identified in the region of grey matter but not around the ependymal canal, with no presence of PCNA-positive cells (Fig 7D, Left). However, in the 7 dpi injured spinal cord, there was a notable increase in the number of cells with nuclear expression of Yap1 around the ependymal canal that also co-express proliferation marker PCNA and a notable reduction in the number of cells with cytoplasmic expression of Yap1 (Fig 7D, Middle). Importantly, under Sirt1-inhibited conditions, the number of cells around the ependymal canal expressing both PCNA and nuclear Yap1 significantly decreased (Fig 7D, Right & 7E) while cells with cytoplasmic Yap1 expression were observed in the grey matter region (Figure 7D, Right). This observation suggests a Sirt1-mediated induction of Yap1 during NPC proliferation. Collectively, these findings suggest that injury-induced Yap1 activation and cytoplasmic-to-nuclear translocation are Sirt1-mediated. According to our hypothesis, this regulation may involve Dnmt1-directed demethylation or hypomethylation of *yap1* promoter.

To prove the affirmative role of Sirt1 and the deeper mechanisms in demethylation/hypomethylation mediated Yap1 activation, we performed target specific MeDIP-sequencing to characterize injury-induced change in DNA methylation in the promoter region of *yap1*. To understand how DNA methylation changes during zebrafish spinal cord regeneration, we collected samples from regenerating spinal cords at two different experimental conditions (DMSO-treated and EX527-treated 7 dpi cord) together with uninjured zebrafish spinal cord. We generated yap1-specific MeDIP-Seq libraries from these samples with a decent amount of CpG coverage (average 11.2× coverage and 62.4% of CpGs covered ≥ 5×).

From the representative tracks and quantification of methylation generated from the alignment sequences, it has been observed that the methylation level of the *yap1* promoter in uninjured spinal cord showed hypermethylation while the promoter became hypomethylated (∼4-fold reduction) compared with the control group (Figure S6A & S6B). Furthermore, EX527 treatment raised the levels of *yap1* promoter methylation by 1.25-folds and 5-folds compared to uninjured and 7 dpi cord, respectively (Figure S6A & S6B). Overall, these results prompted that upon injury the promoter of *yap1* became hypomethylated which leads to the transcriptional activation of *yap1* while Sirt1-inhibition leads to the hypermethylation of *yap1* promoter region which caused reduced transcriptional activity in 7 dpi spinal cord.

### 3.9. Sirt1 promotes expression of *ctgfa* through induction of Yap1 mediated HIPPO signalling pathway

To prove that Sirt1 activates Yap1 and its nuclear translocation and subsequent increased expression of *ctgfa,* we observed the endogenous level of *ctgfa* during regeneration time course in both DMSO-treated control and EX527-treated fish by means of RT-PCR method. The mRNA expression pattern during regeneration time course showed gradual increase with a high expression level at 7 dpi and 15 dpi when compared to the uninjured control (Fig. 8A) which is in line with the previous observation by Mokalled et al,2016(7). But in case of Sirt1-inhibited condition at 7 dpi, the expression has been reduced as observed from the RT-PCR analysis (Fig. 8A). To further elucidate the role of Sirt1-mediated post-translational modification of Yap1 in inducing *ctgfa* transcription, we used a transgenic reporter zebrafish with a 5.5 kb genomic sequence upstream of the *ctgfa* translational start site fused to a EGFP reporter cassette. In case of DMSO-treated control condition, analysis of *ctgfa*:EGFP expression and GFAP expression, a marker of glial cells in the CNS revealed a higher expression of Ctgfa at the site of injury at 7 dpi and overlapping of *ctgfa*:EGFP and GFAP within a subpopulation of glial cells at the injury site (Fig. 8B) which indicates the bridging cells, as also described by Mokalled et al.(7). On the contrary, fish from Sirt1-inhibited condition showed significantly reduced expression of ctgfa at the injury site and no bridging cells with co-expression of both *ctgfa*:EGFP and GFAP has been observed indicating that functional inhibition of Sirt1 during early time point of regeneration directs the reduced expression of Ctgfa (Fig. 8B) and as per our experimental observations, this regulation is controlled by Sirt1-mediated activation of Yap1.

**Figure 8.**
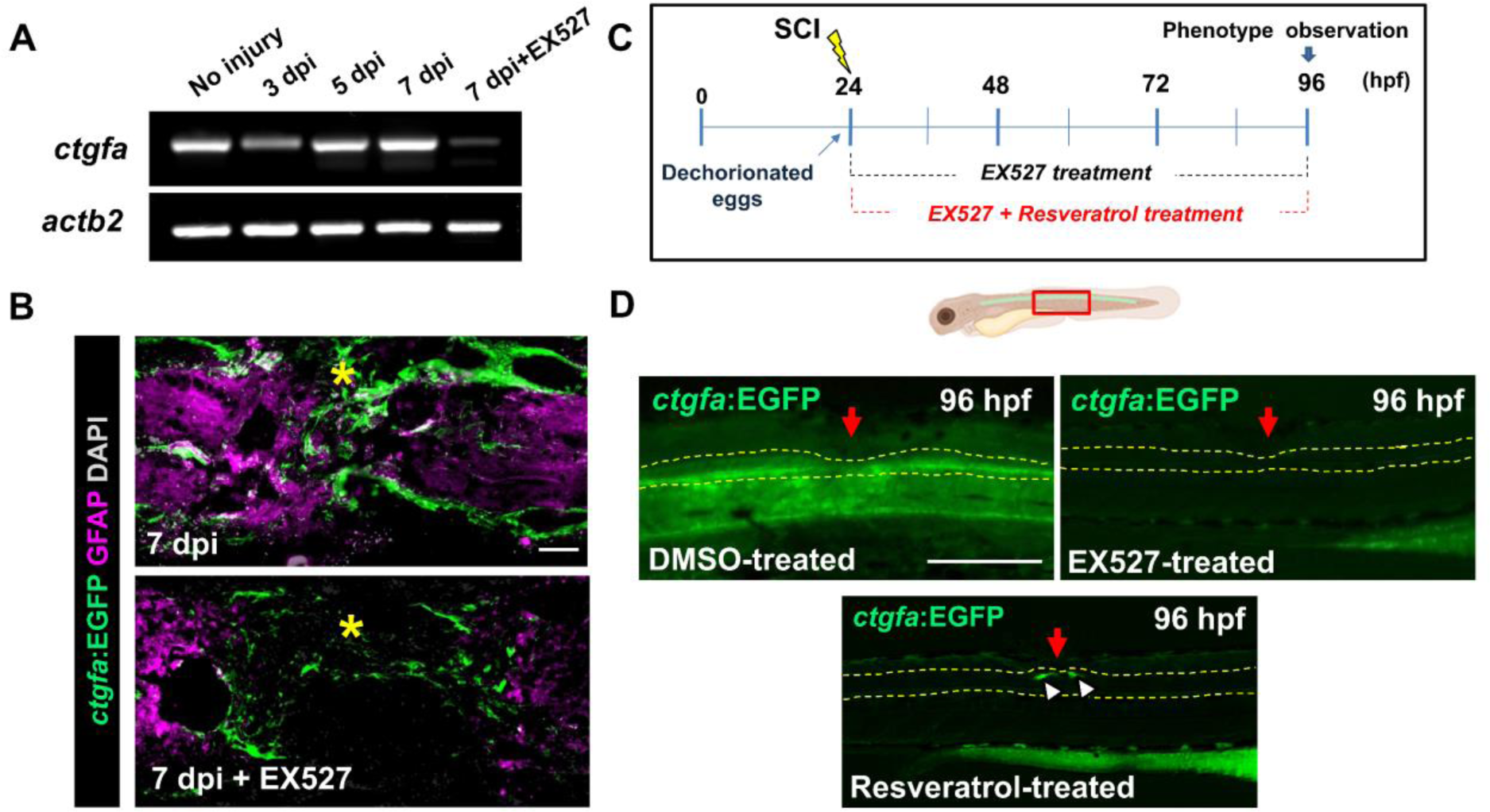
Legend: Sirt1-mediated activation of *ctgfa* during spinal cord regeneration. (A) RT-PCR analysis of endogenous ctgfa expression in during regeneration and Sirt1-inhibited condition. (B) Immunohistochemical staining of 7 dpi injured spinal cord (Transverse Section) in uninjured (left), DMSO-treated (middle) and EX527-treated (right) experimental groups showing co-expression of *ctgfa*:EGFA and GFAP. Yellow star indicates the injury epicentre. Magnification, 20x; Scale bar, 100µm. (C) Scheme of experimental conditions using *ctgfa:*EGFP embryos after SCI in DMSO-treated control, EX527-treated and Resveratrol-treated condition. (D) Whole-mount immunohistochemical staining of *ctgfa*:EGFP embryos at 96 hpf stage in different experimental conditions. Yellow dotted lines demarcate the spinal cord and the red arrow indicates the injury point. Magnification, 10x, scale bar 100 µm.

We further observed the role of Sirt1 on ctgfa expression in embryonic condition using *ctgfa*:EGFP embryo. Spinal cord injury was given at 24 hpf embryo after dechorionation and treated with DMSO and EX527 for 3 days up to 96 hpf (Figure 8C). At 96 hpf, increased expression of *ctgfa* was observed in the spinal cord surrounding the injury epicentre while, remarkably, EX527-treated embryos showed completely reduced *ctgfa*:EGFP expression around the injury epicentre (Figure 8D). Embryos treated with combination of EX527 and Resveratrol showed comparatively increased *ctgfa*:EGFP expression at the injury epicentre (Figure 8D) which suggest that resveratrol can rescue the inhibitory condition of EX527 and induce Sirt1 activity. Altogether, these results support our preceding experiments that Sirt1 can regulate *ctgfa* expression by modulating HIPPO Pathway not only in adult stage but also during embryonic stage of zebrafish.

## 4. Discussion

Epigenetic modifications serve as mechanisms employed by cells to modulate gene expression in response to injury. A recent investigation demonstrated that inhibiting histone deacetylase (HDAC) influenced neuroepithelial (NE) proliferation under physiological conditions in the adult mice optic tectum (61). Furthermore, histone acetylation has been identified as a regulatory factor in determining the regenerative capacity of both mammalian and zebrafish retina (62,63), as well as the zebrafish optic tectum (64). In the present study, we demonstrate that the epigenetic modulator Sirt1 is functionally involved in the proliferation of NPC in the regenerating spinal cord of zebrafish. We observed that that increased Sirt1 activity is associated with bridging activity of neural progenitor cells along with axonal regrowth and demonstrate that inhibition of Sirt1 expression in zebrafish during spinal cord regeneration not only prevents NPC proliferation but also impedes glial bridging and axonal regeneration. Moreover, we demonstrate that NPC proliferation during regeneration is controlled by the epigenetic modulation of HIPPO signalling pathway, in particular by deacetylation-mediated inactivation of Dnmt1 and subsequent demethylation of *yap1* that depend on Sirt1 activation. These findings reveal that Sirt1 activity is essential to both NPC proliferation and axonal regeneration.

The expression of Sirt1 was specifically upregulated in early regenerative time points in zebrafish spinal cord. Previously it has been reported that in zebrafish Sirt1 is associated with the aging, neuroinflammation, microglia activation in zebrafish but so far, the epigenetic role of Sirt1 has not been studied during spinal cord regeneration in zebrafish. In case of mammals, SIRT1 levels gradually decreased in spinal cord until the fourth week after SCI in rat (65), while, on the contrary, another study observed that SIRT1 began to decline 4 hours after SCI, and the minimum levels were observed at 8h post-injury (hpi). SIRT1 expression returned to the normal level by the third day (20). We observed increased expression of *sirt1* in both RT-PCR and indirect ELISA at around 7 dpi which coincides with proliferation pattern of injury-induced NPCs during regeneration. Thus, we aimed to observe the implication of Sirt1 in injury-induced NPC proliferation and our loss-of-function studies confirmed reducing NPC proliferation after Sirt1 inhibition and rescuing the loss-of-function effect after Sirt1 overexpression. Similar to our findings others also reported that Sirt1 regulates glial progenitor proliferation and regeneration in white matter after neonatal brain injury in mice (32).Therefore we establish Sirt1 as an essential component of the regulatory mechanism that controls NPCs proliferation during spinal cord regeneration.

SIRT1 was also shown to be expressed in most of the SOX2-expressing proliferating Oligodendrocyte precursor cells (OPCs) in subventricular zone of mice brain (66). In case of zebrafish spinal cord, we observed by immunohistochemical analysis that all Sox2^+^ NPCs present around the ependymal canal were Sirt1-positive and thereby indicating injury-induced Sirt1 activation in the NPCs. Direct functional evidences were obtained both *in vitro* and *in vivo* from previous reports, demonstrating that after chronic neonatal hypoxic condition, increased cytoplasmic SIRT1 in OPCs strongly correlates with their proliferative state, suggesting that the proliferative response of OPCs is mainly regulated by SIRT1. Also, downregulation of SIRT1 protein expression in cultured OPCs, as well as excision of the *Sirt1* gene in Sirt1-cKO mice severely reduced OPC expansion after hypoxia. Similar result was obtained in our study showing reduced NPC proliferation upon Sirt1-inhibition and together these results suggested that injury-induced Sirt1-activation in NPCs is associated with NPC proliferation. Thus, it will be important to address in which way Sirt1 as an epigenetic modulator control the regulatory mechanism of NPC proliferation during spinal cord regeneration and whether it also has some positive role in other regenerative events other than NPC proliferation such as glial bridging and regenerative axonogenesis leading to gain functional recovery after SCI. To address the question, we observed the loss-of-function effect of Sirt1 on glial bridging and reported a significant reduction in glial bridge formation upon sirt1-inhibition at 21 dpi. Previously no reports have been found showing the association of any epigenetic factor with induction of NPC proliferation and also in glial bridge formation. Our study suggest that Sirt1 is critically required for both the induction of NPC proliferation along with the changes in morphology of glial cells and glial bridge formation during spinal cord regeneration.

SIRT1 promotes axon outgrowth by deacetylating and activating Akt and ultimately inactivating GSK3 in cultured rat embryonic hippocampal neurons (33). Similar result was also reported by Romeo-Guitart et al (2019), showing that SIRT1 overexpression on spinal motoneurons facilitates a growth-competent state improving motor axonal regeneration in mice (67). Several other studies also reported that SIRT1 plays a central role in axonogenesis and optic nerve regeneration (34,68,69). In support with the previous observations here we report that Sirt1-inhibition leads to disrupted axonal regrowth and locomotor behavioural analysis supports the observation suggesting that Sirt1 induce axonal regrowth after injury and also facilitate to recover functional behaviours.

During zebrafish spinal cord regeneration, NPC proliferation is accompanied by the increased expression of pluripotency and other regeneration-associated genes. Previously studied somatic cell reprogramming events found strong correlations between the increased expression of pluripotency factors and decreases in their promoter methylation levels (70–72). Likewise, we hypothesized that the regulation of DNA methylation would accompany the regenerative events in the injured spinal cord. Powell et al(2013) reported that during Müller glia-derived progenitor cells (MGPC) formation (0–2 dpi) in case of retinal regeneration in zebrafish, DNA demethylation predominates at early times, whereas levels of de novo methylation increase at later times which indicate a noted correlation between promoter DNA methylation and injury-induced gene induction (46). They also observed that the promoters of pluripotency factors were hypomethylated in quiescent MG and remained unchanged in MGPCs. Indeed, we found that during early regenerative time points (3 dpi and 7 dpi), zebrafish genome undergoes dynamic changes in global DNA methylation showing a significant decrease in global 5-mC percentage during early time points which increased at later time point (15 dpi). Here, we observed induction in the global 5mC percentage upon Sirt1 inhibition at the early time points of regeneration (3 dpi and 7 dpi). From these observations, we hypothesize that underlined mechanisms must also be at play because some of the best-studied regeneration-associated genes do not undergo changes in DNA methylation. Other additional factors such as availability of transcription factors and other epigenetic changes may directly regulate these genes. Thus, changes in DNA methylation may be necessary for progenitor cell reprogramming, but may not be sufficient.

Histone modifications are one of the possible mechanisms that progenitors use to maintain its self-renewal capabilities and to alter their fate choices between neurogenesis and gliogenesis (73,74). Recent study has shown that histone deacetylase (HDAC) inhibition upregulates Notch signalling and reduce cell proliferation in the optic tectum of adult zebrafish (61). Moreover, it was reported that HDAC inhibition after stab wound injury suppressed RG proliferation but promoted neuronal differentiation in zebrafish optic tectum (64). Mitra et al (2018) also reported the similar observation that hdac1 is essential for MGPC proliferation during zebrafish retinal regeneration (63). In case of zebrafish spinal cord, a recent study observed that Hdac1 induce newborn motor neuron formation by activating ERG proliferation (75). Thus, it has been obvious that histone acetylation and deacetylation play an essential role in injury-induced regenerative responses in zebrafish CNS. In that perspective, the role of other histone modifiers remains less explored. Likewise, in our study we hypothesized a positive association between sirt1-mediated deacetylation and NPC proliferation and subsequent axonal regrowth. For that purpose, we observed the pan-acetylation level in the nuclear extract of experimental zebrafish which reflected a gradual decrease in global acetylation level in the early time points (3 dpi and 7 dpi) while Sirt1-inhibition slightly induced the acetylation level (1-fold) suggesting that at the global level acetylation is not completely regulated by Sirt1 alone. To our surprise, EX-527 mediated Sirt1 inhibition did not significantly increase H4 acetylation rather than induce H3 acetylation level. We conclude that Sirt1 might not be the primary deacetylase of acetylated H4 and that global H4 acetylation levels may not always be accurate indicators of cellular Sirt1 deacetylase activity.

DNA methylation is catalyzed by a family of DNA methyltransferases (DNMTs), including DNMT1, DNMT3A, and DNMT3B (76). While DNMT1 primarily detects hemimethylated DNA and maintains DNA methylation patterns in dividing somatic cells, DNMT3A and DNMT3B are de novo methyltransferases that covalently add a methyl group to the C5 of cytosine in CpG dinucleotides. (77). However, the classification is not tightly restricted. Based on sequence similarity, the existence of mammalian-like Dnmt1 and Dnmt3 proteins in zebrafish implies that pathways linked to maintenance and de novo methylation are broadly conserved across species (50). In zebrafish, Dnmt1 is primarily responsible for maintenance of methylation while all the members of Dnmt3 group are associated with de novo methylation (78). Limited evidence suggests the presence of de novo activity of DNMT1 (79–81), but the overall evidence was not immediately clear. A recent study unveiled that DNA methylation, which generally leads to gene silencing, also contributes to regenerative responses of conditioned DRG neurons (82). In a sciatic nerve damage model, peripheral nerve regeneration is decreased by treatment with RG108, a direct DNMT inhibitor. It has been reported that DNMT1 has a crucial effect on global genomic methylation. For example, the inactivation of DNMT1 causes DNA demethylation, and homozygous null deletions of DNMT1 result in an 80% genomic loss of DNA methylation in mouse (83,84). In human cells, DNMT1 transcription steadily declines throughout the aging process (85). These findings suggest that in vertebrates, reduced genome-wide methylation during aging can be attributed to a decreased abundance of DNMT1.Likewise, we observed a significant decrease in the mRNA expression of *dnmt1* and *dnmt2* but no significant changes in the other *dnmt3* members at 7 dpi. Also, a gradual decline in global 5-mC percentage which we can correlate to the similar kind of decreased expression of both *dnmt1* transcript and protein during the early regenerative time points (3-7 dpi). Moreover, upon Sirt1 inhibition, both the global methylation level as well as expression of *dnmt1* transcript and protein level have been increased significantly throughout the regeneration time course in comparison to the condition when Sirt1 is functionally active. Therefore, we could assume that Sirt1 modulates the global DNA methylation by directing the *de novo* activity of Dnmt1 rather than its maintenance role. In our study, we also observed a notable alteration in the expression of *dnmt2* during regeneration, as determined by RT-PCR. Considering that *dnmt2* (*tRNA aspartic acid methyltransferase 1*) exhibits strong methyltransferase activity on tRNA^ASP^ (86,87) and minimal levels of Dnmt2-dependent DNA methylation (88), further investigation is necessary to fully understand the role of *dnmt2* in spinal cord regeneration in zebrafish.

The mechanisms by which DNMT1 controls gene expression have been much explored. However, the mechanisms governing the abundance and activity of DNMT1 remain less understood, although emerging evidence suggests a potential role of DNMT1 behind the post-translational modifications. A few studies demonstrated that acetylation of DNMT1 may associate with changes in catalytic activity, DNA binding activity, and/or stability (89,90). Peng et al(2011) reported SIRT1 mediated deacetylation of DNMT1 which changed its activity to silence downstream gene expression, and its ability to regulate the cell cycle (91). But in the context of zebrafish spinal cord regeneration the mechanism of Dnmt1 activity in regulating injury-induced regenerative events is still an unexplored area. Here, we observed the reduced acetylation of H3 and H4 protein in the nuclear extract after immunoprecipitation using Dnmt1 antibody mostly around 7 dpi when NPC proliferation is in peak. Upon Sirt1 inhibition, the acetylated H3 level increased significantly in our study rather than H4, which is at the same time consistent with our previous finding where Sirt1 inhibition increased the global Acetylated H3 level rather than H4. These findings reveal a novel cross talk between two important epigenetic effectors, Dnmt1 and Sirt1 during zebrafish spinal cord regeneration.

Earlier reports have indicated that NPCs in the zebrafish spinal cord, specifically ERGs, undergo dedifferentiation and elongate their processes in response to injury. In conjunction with regrowing axons, these cells actively contribute to the formation of a glial bridge, thereby facilitating the reconnection of the severed spinal cord (7,92,93). These processes are facilitated by several signalling pathways, such as Wnt/β-catenin, Fgf, Shh, while being inhibited by Notch signalling. To initiate the bridging process, ventral ERGs undergo an epithelial-to-mesenchymal transition (EMT). which is a prevalent characteristic observed in glial cells responding to injury and is associated with stem cell activation, increased cellular plasticity, and tissue remodelling (55,94,95). This process of Glial EMT is indispensable sufficient to initiate glial bridging. This phenomenon is closely associated with the Yap1-Ctgfa signalling pathway, also recognized as the HIPPO signalling pathway (59).

Earlier studies have consistently revealed a direct correlation between the expression levels of YAP and SIRT1 through the utilization of knockout and overexpressing mice (96). Mao et al. demonstrated a significant up-regulation of SIRT1 expression in hepatocellular carcinoma samples, wherein the SIRT1 mRNA level showed a positive correlation with *Ctgf*, a target gene of YAP(97). Additionally, it was observed in hepatocellular carcinoma cells that SIRT1 deacetylated the YAP2 protein. The SIRT1-mediated deacetylation, in turn, heightened the association between YAP2 and TEAD4, resulting in enhanced the transcriptional activation of YAP2/TEAD4 and increased cell proliferation in HCC cells(97). However, the exact relationship between YAP and SIRT1, specifically whether YAP acts as a direct target of SIRT1 or functions as a downstream target, has yet to be conclusively clarified. We observed an augmented Yap1 activity in the tissue lysate over the entire regeneration time course, particularly at 7 dpi spinal cord. A similar trend was noted in Yap1 activity within the nuclear extract, with lower nuclear Yap1 levels at the early time points (3 and 5 dpi) compared to 7 dpi spinal cord, while the cytoplasmic Yap1 level was low at that time point where NPC proliferation was higher. These findings suggested a cytoplasmic-to-nuclear translocation of Yap1 during NPC proliferation. Moreover, we observed a significant reduction in nuclear Yap1 levels upon Sirt1 inhibition, whereas cytoplasmic Yap1 levels remained unaffected. Similar kind of observation was also noted by immunohistochemical analysis where Yap1 was expressed in the cytoplasm but upon injury, the cells undergo EMT and located around the ependymal canal and showed cytoplasmic-to-nuclear translocation of Yap1 and subsequent proliferation of the cells which were validated by the proliferation marker PCNA. But upon Sirt1-inhibition, cytoplasmic retention of Yap1 has been observed leading to reduced proliferation. It implies that Sirt1 plays a pivotal role in directing the activation and nuclear translocation of Yap1. The promoter region of *yap1* gene contains many CpG and non-CpG islands which are the most promising targets for epigenetic regulation. To observe the deeper mechanism behind the Sirt1-induced change in methylation pattern of *yap1* during regeneration, we performed target-specific MeDIP-sequencing and observed that the promoter region of *yap1* gene become hypomethylated post-injury at 7 dpi and reduced 1.25-folds in comparison to uninjured animals (Figure S6). However, hypermethylation of the promoter region was observed in case of Sirt1-inibited condition at 7 dpi which has been increased around 5-folds compared to DMSO-treated 7 dpi cord (Figure S6). One recent study reported the role of DNMTs in hypermethylation the promoter region of YAP1 in breast cancer cells (98). Therefore, we can correlate the expression pattern of *dnmt1* with the methylation pattern of *yap1* promoter and it strongly suggests that during spinal cord regeneration, increased Sirt1 level causes deacetylation of Dnmt1 leading to its inactivation and at that timepoint *yap1* promoter becomes hypomethylated due to reduced Dnmt1 activity. When Sirt1 was functionally inhibited at 7 dpi regenerating cord, Dnmt1 became more activated in comparison to uninjured condition where Sirt1 level was low. This causes hypermethylation of the *yap1* promoter and subsequent transcriptional inactivity of *yap1* gene. Also, remarkably, in contrast with non-CpG methylation, we found that injury-induced alterations in CpG methylation in the *yap1* promoter were minimal. This observation is in line with the study conducted by Reverdatto et al. (2022) which showed that the change in the methylation of non-CpG sites were comparatively higher in comparison to CpG sites in CNS of Xenopus (99).

Altogether, our study suggests that Sirt1-mediated change in the methylation of *yap1* promoter through modulation of Dnmt1 acetylation is the principal machinery behind the injury-induced inactivation of HIPPO pathway in zebrafish spinal cord regeneration. Additionally, we also found a decrease in *ctgfa*:EGFP expression under the conditions where Sirt1 was inhibited in both embryonic and adult stage. It is well known that Ctgfa is a pro-regenerative factor which induce the glial bridge formation and axonal regrowth through Yap1-mediated modulation of HIPPO pathway. Here we elucidate that Sirt1 is the key epigenetic factor behind the activation of *ctgfa* upon injury through Dnmt1-mediated methylation of Yap1, a major intermediate factor of HIPPO pathway.

In conclusion, we show that in uninjured zebrafish spinal cord characterized by a low level of Sirt1, Dnmt1, being acetylated maintains an active status and by observing the Yap1 expression it can be assumed that activated Dnmt1 methylate *yap1* promoter leading to inactivation of Yap1, thereby activates the HIPPO signalling pathway. Following injury, activated Sirt1 deacetylates Dnmt1, resulting in the repression of both its methyltransferase activity and transcription repression capability and subsequently impairs its capacity to inactivate Yap1. Upon demethylation, Yap1 becomes activated, experiencing cytoplasmic-to-nuclear translocation, and subsequently instigates the transcription of the pro-regenerative factor, *ctgfa*. Our investigation unveils a novel insight into the Sirt1-mediated post-translational regulation of Yap1, establishing a previously undisclosed Sirt1-Dnmt1-Yap1 cross-talk which provides a deeper understanding of the epigenetic regulatory mechanisms governing injury-induced NPCs proliferation, glial bridging, and subsequent axonal regrowth in the zebrafish spinal cord.

## Author Contributions

SG and SPH developed concept and designed experiments. SG conducted experiments, collected data, and performed analysis. SG and SPH wrote manuscript. SPH supervised all aspects of the project.

## Acknowledgments

We thank University Grant Commission, Govt. of India for their financial support through the UGC START-UP GRANT [F.30-595/2021(BSR) GEN-31] and Department of Science and Technology-Science and Engineering Research Board (DST-SERB), Govt. of India for their financial assistance through the Core Research Grant (CRG/2022/001663) to SPH. SG is a recipient of the Senior Research Fellowship (SRF) award [09/028(1145)-2020-EMR-I] from the Council of Scientific & Industrial Research (CSIR), Govt. of India.

## Supplementary Materials

**Supplementary Figure S1.**
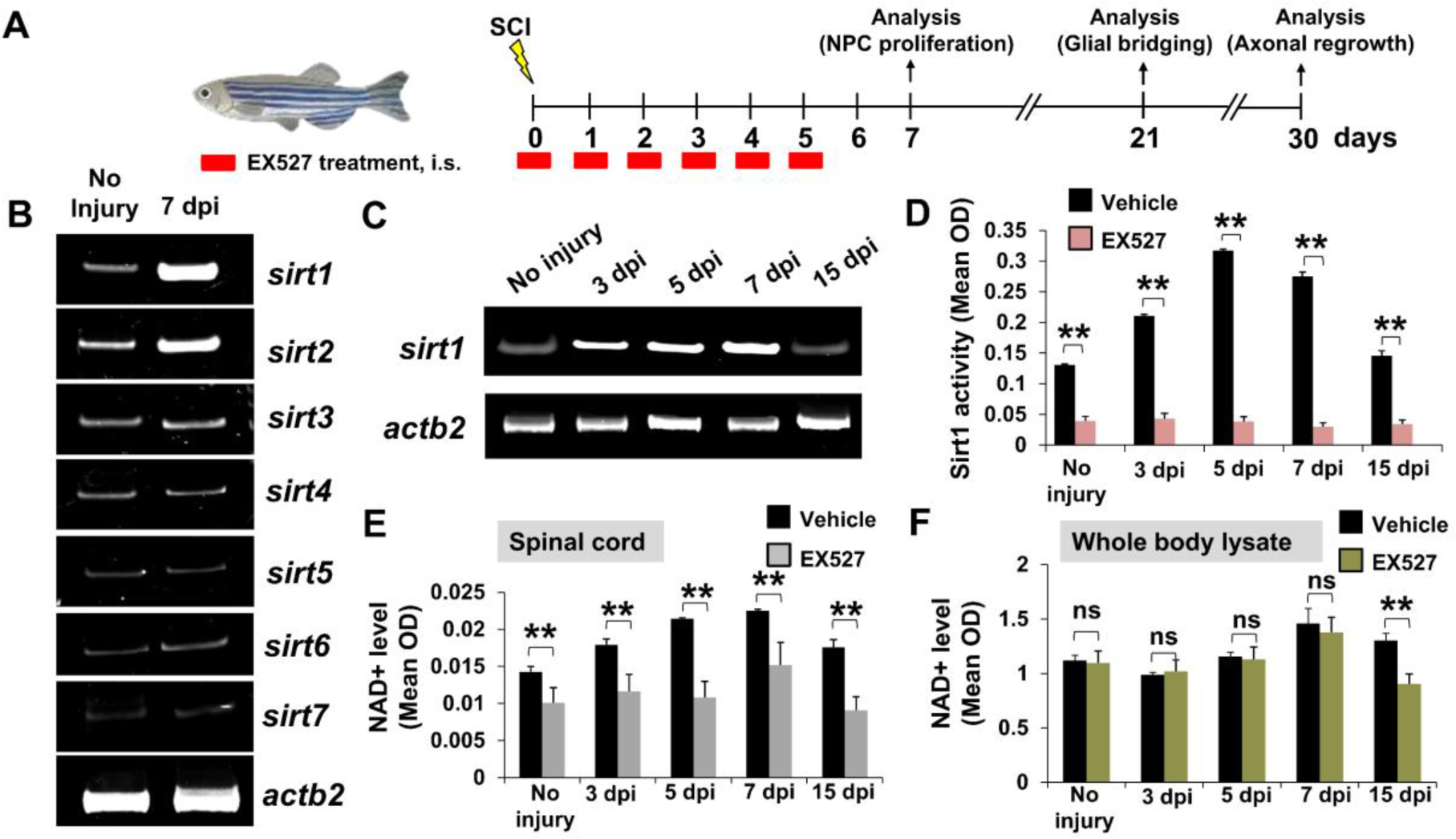
**Legend:** (A) Experimental intervention involving EX527 treatment. i.s. intra-spinal injection. (B) RT-PCR analysis of expression of *sirtuin* members in regenerating spinal cord after 7 dpi; (C) RT-PCR analysis of expression of *sirtuin1* during regeneration time course in zebrafish spinal cord (D) Quantitative analysis of Sirt1 protein levels in different time points by indirect ELISA method in DMSO-treated control and EX527-treated experimental group of zebrafish after SCI. (D) Quantitative analysis of time course specific NAD^+^ level in spinal cord tissue by cyclic NAD assay in DMSO-treated control and EX527-treated experimental group of zebrafish after SCI. (E) Quantitative analysis of time course specific NAD^+^ level in whole body lysate in DMSO-treated control and EX527-treated experimental group of zebrafish after SCI. Data represented as Mean±;SEM, n=6; \**p* < 0.05; \*\**p* < 0.01; ns, Non-significant; Mann-Whitney U test.

**Supplementary Figure S2.**
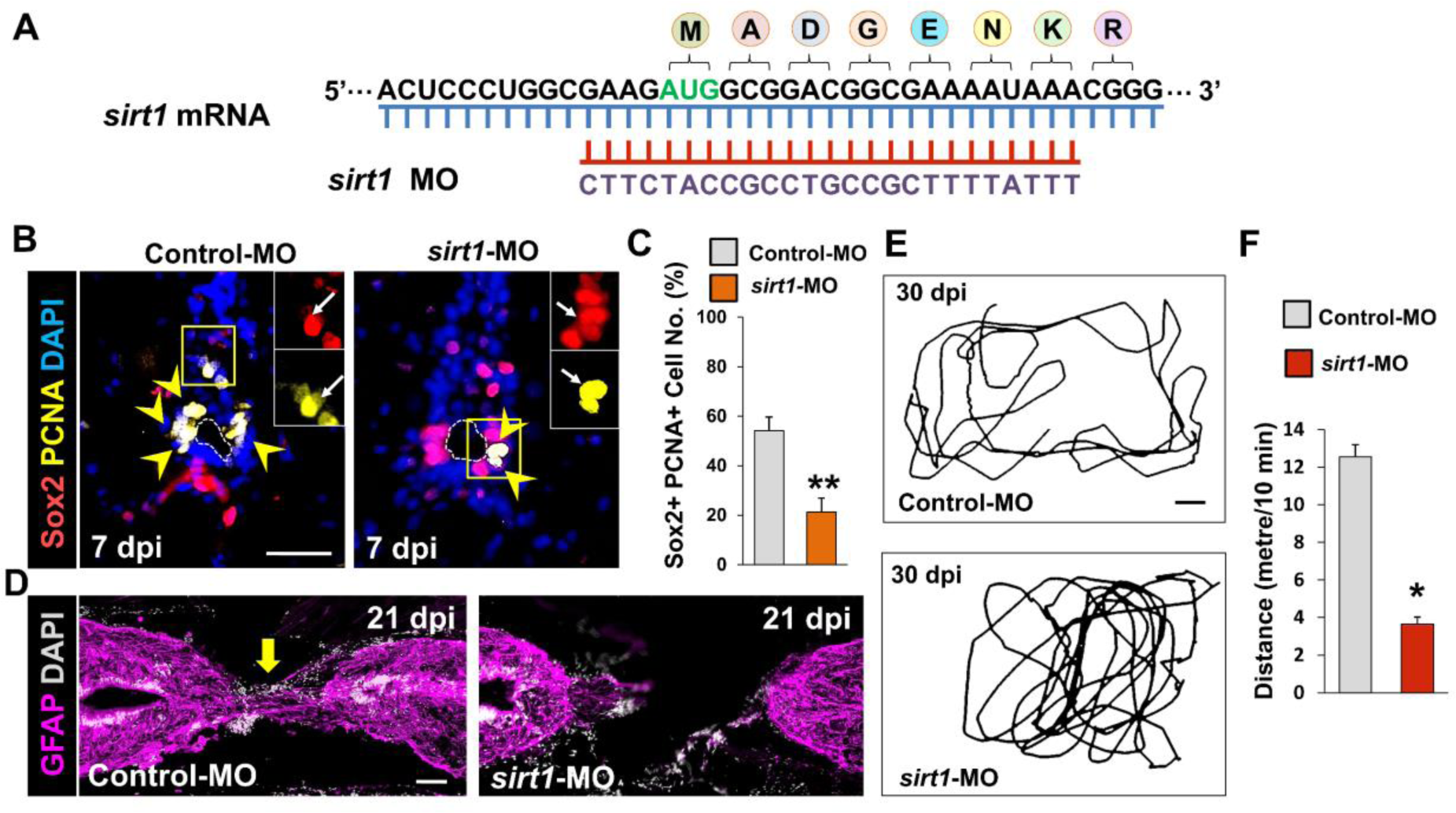
**Legend:** (A) Schematic diagram to depict base-pairings between *sirt1* mRNA and antisense morpholino oligonucleotides (MO) of *sirt1* (*sirt1*-MO); (B) Immunohistochemistry of transverse section of zebrafish spinal cord showing Proliferating neural progenitor cells (co-expression of Sox2 & PCNA, marked by yellow arrowheads) after 7 dpi in both Control-MO treated (left) and *sirt1*-MO treated (right) condition. The white-dotted line marked the ependymal canal. Magnification, 20x; Insets show single-channel slices of the demarcated region; (C) Quantification of (B) (N=5); (D) Immunohistochemical staining of spinal cord sections after 21 dpi from Control-MO treated (left) and *sirt1*-MO treated (right) fish showing glial bridge formation. Yellow arrow indicates glial bridging; (E) Swim tracking of individual animals after 30 dpi from EX527-treated (Upper) and Resveratrol-treated (Lower) experimental groups (N=5); (F) Quantification of total distance travelled in (E). Data represented as Mean±SEM, n=5; \**p* < 0.05; \*\**p* < 0.01; ns, Non-significant; Mann-Whitney U test. Scale bars, 50 µm (B, D) and 0.1 metre (E).

**Supplementary Figure S3.**
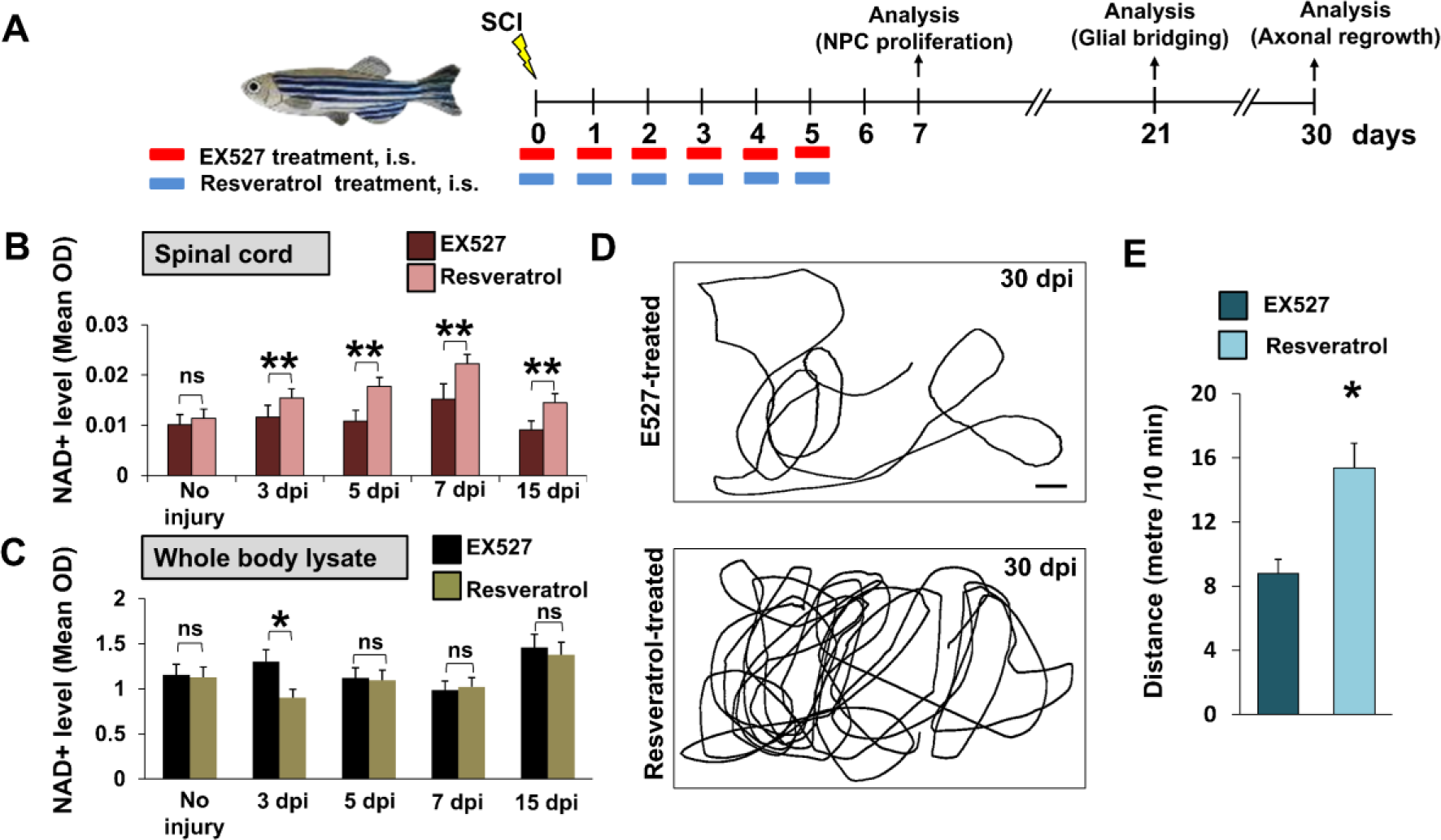
**Legend:** (A) Experimental intervention involving EX527 and Resveratrol treatment. i.s. intra-spinal injection. (B) Quantitative analysis of time course specific NAD^+^ level in spinal cord tissue by cyclic NAD assay in EX527-treated control and Resveratrol-treated experimental group of zebrafish after SCI. (N=5). (C) Quantitative analysis of time course specific NAD^+^ level in whole body lysate in in EX527-treated control and Resveratrol-treated experimental group of zebrafish after SCI (N=5). (D) Swim tracking of individual animals after 30 dpi from EX527-treated (Upper) and Resveratrol-treated (Lower) experimental groups (N=8). (E) Quantification of regeneration in (D). Data represented as Mean±SEM, n=5; \**p* < 0.05; \*\**p* < 0.01; ns, Non-significant; Mann-Whitney U test. Scale bar, 0.1 metre (D).

**Supplementary Figure S4.**
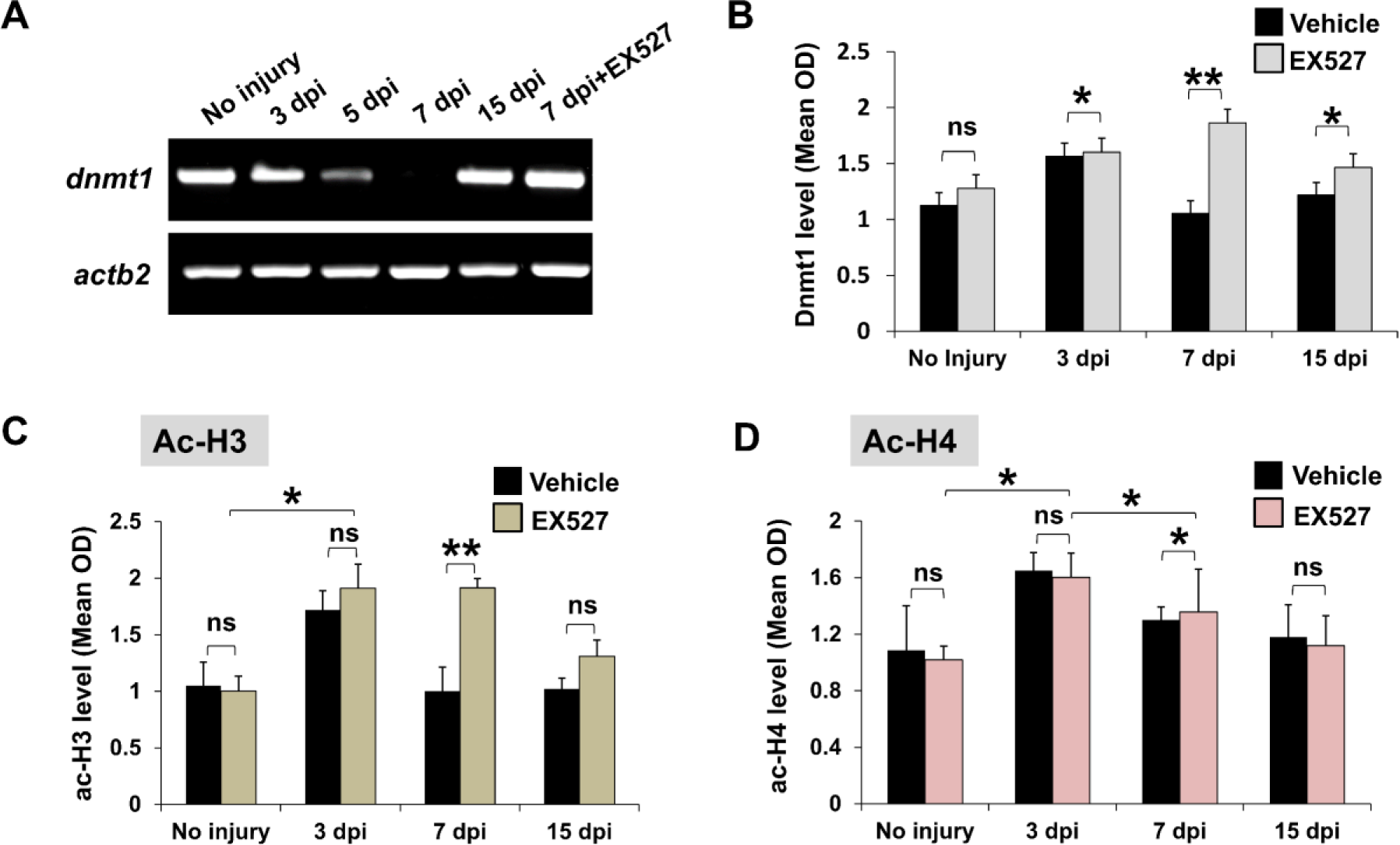
**Legend: (**A) RT-PCR analysis of expression of *dnmt1* in regenerating spinal cord after 7 dpi; (B) Quantitative analysis of Dnmt1 protein level in different time points by indirect ELISA method in DMSO-treated and EX527-treated experimental groups; (C, D) Quantitative analysis of Dnmt1 specific acetylated-H3 and acetylated-H4 level in nuclear extract from spinal cord tissues method in DMSO-treated and EX527-treated experimental groups by immunoprecipitation followed by indirect ELISA method. Data represented as Mean±SEM, \**p* < 0.05; \*\**p* < 0.01; ns, Non-significant; Mann-Whitney U test.

**Supplementary Figure S5.**
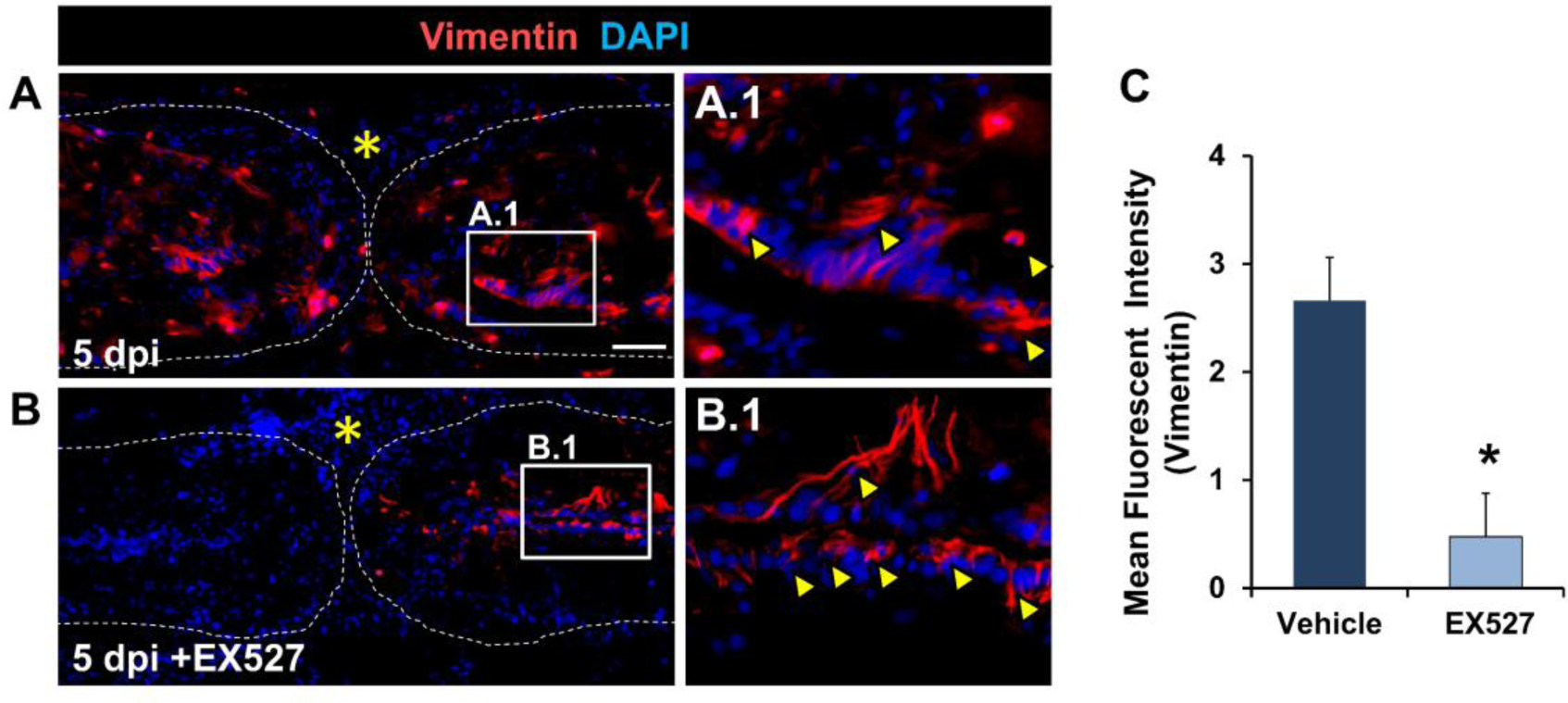
Legend: sirt1-mediated regulation of epithelial-to-mesenchymal transition in spinal cord regeneration. (A) Immunohistochemical staining of 5 dpi injured spinal cord (longitudinal section) in DMSO-treated (upper) and EX527-treated (lower) experimental groups showing expression of vimentin. The white-dotted line marked the spinal cord outline. Magnification, 20x; Insets show higher magnification of the demarcated region; White arrow marked the Vim-positive cells; Yellow asterisk indicates injury epicentre; Scale bar, 50µm; (B) Quantification of (A). Data represented as Mean±SEM, \**p* < 0.05; Mann-Whitney U test.

**Supplementary Figure S6.**
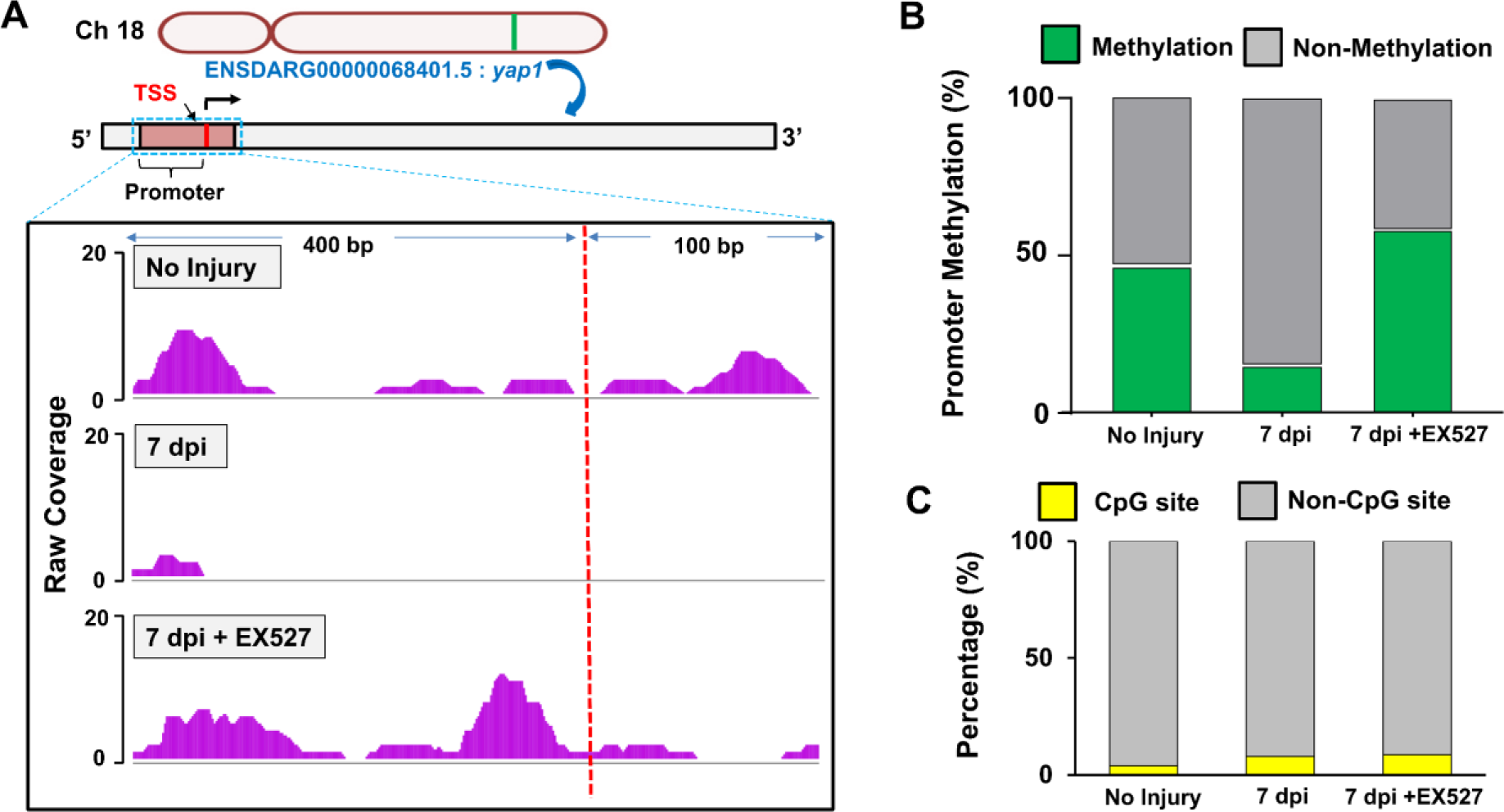
**Legend:** (A) Chromosomal location of *yap1* gene in zebrafish genome and representative screenshots from sequence tracks at zebrafish *yap1* promoter region (Covering -400 bp upstream and 100 bp downstream of Transcription Start Site (TSS, marked by red dotted line) from alignment files of MeDIP-seq samples: (top) Uninjured spinal cord, (middle) DMSO-treated 7 dpi cord, and (bottom) EX527-treated 7 dpi cord. Coverage graphs on top of their respective tracks. (B) Quantification of percentage of promoter methylation in the experimental groups including uninjured group of animals. (C) Quantification of methylated CpG sites and non-CpG sites across the *yap1* promoter region in different experimental groups.

## Key Resource Table

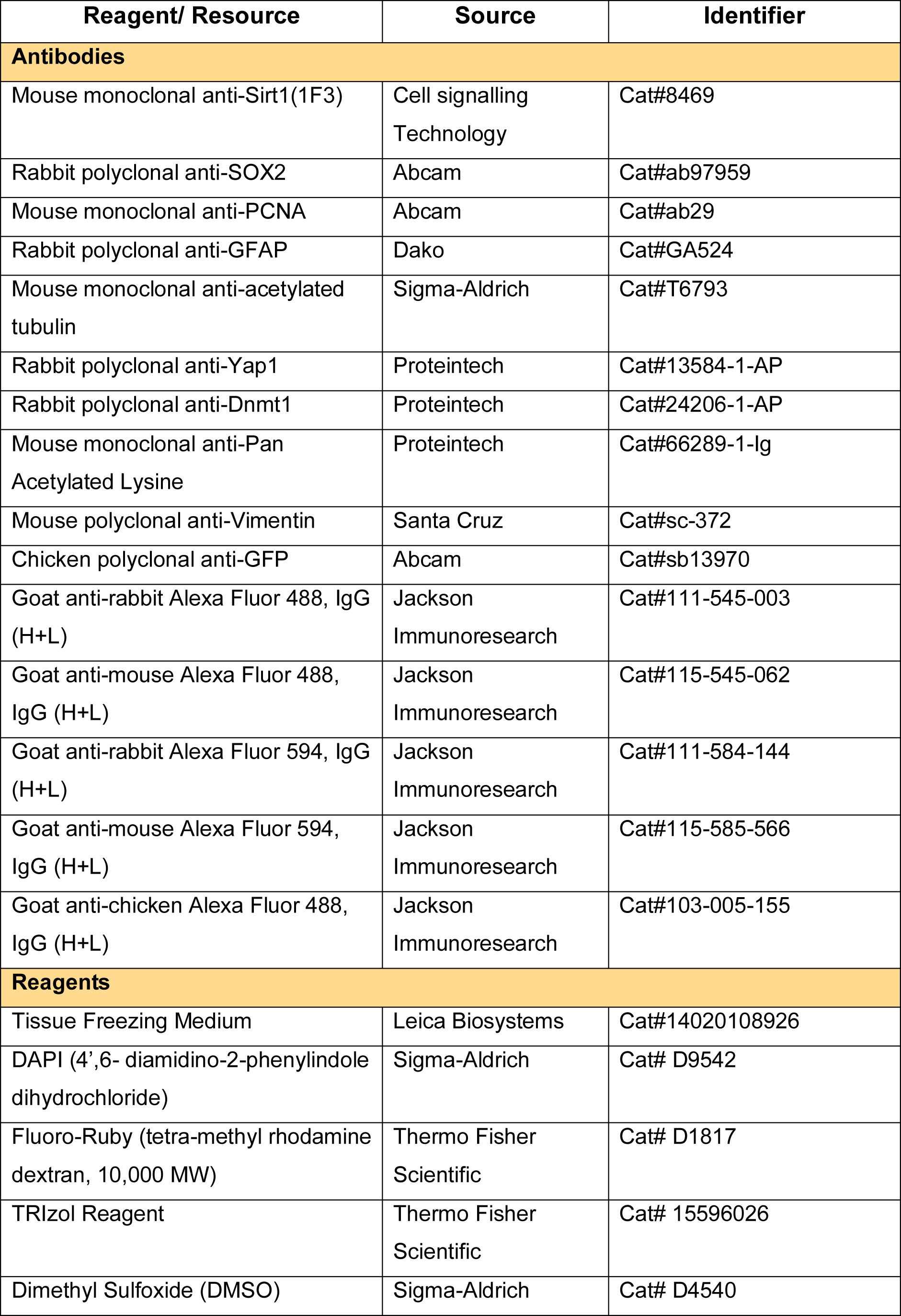
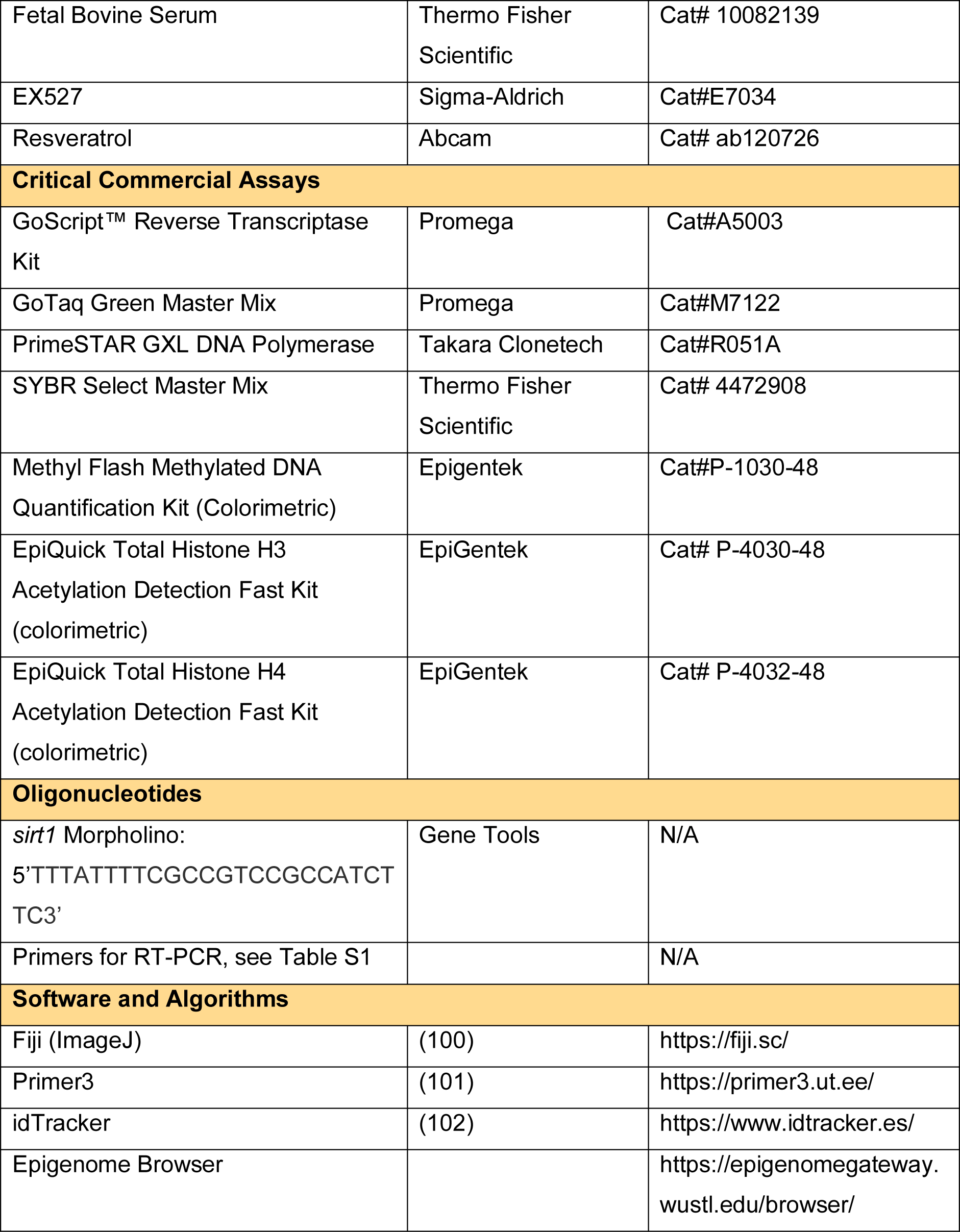

---

**Table S1:**
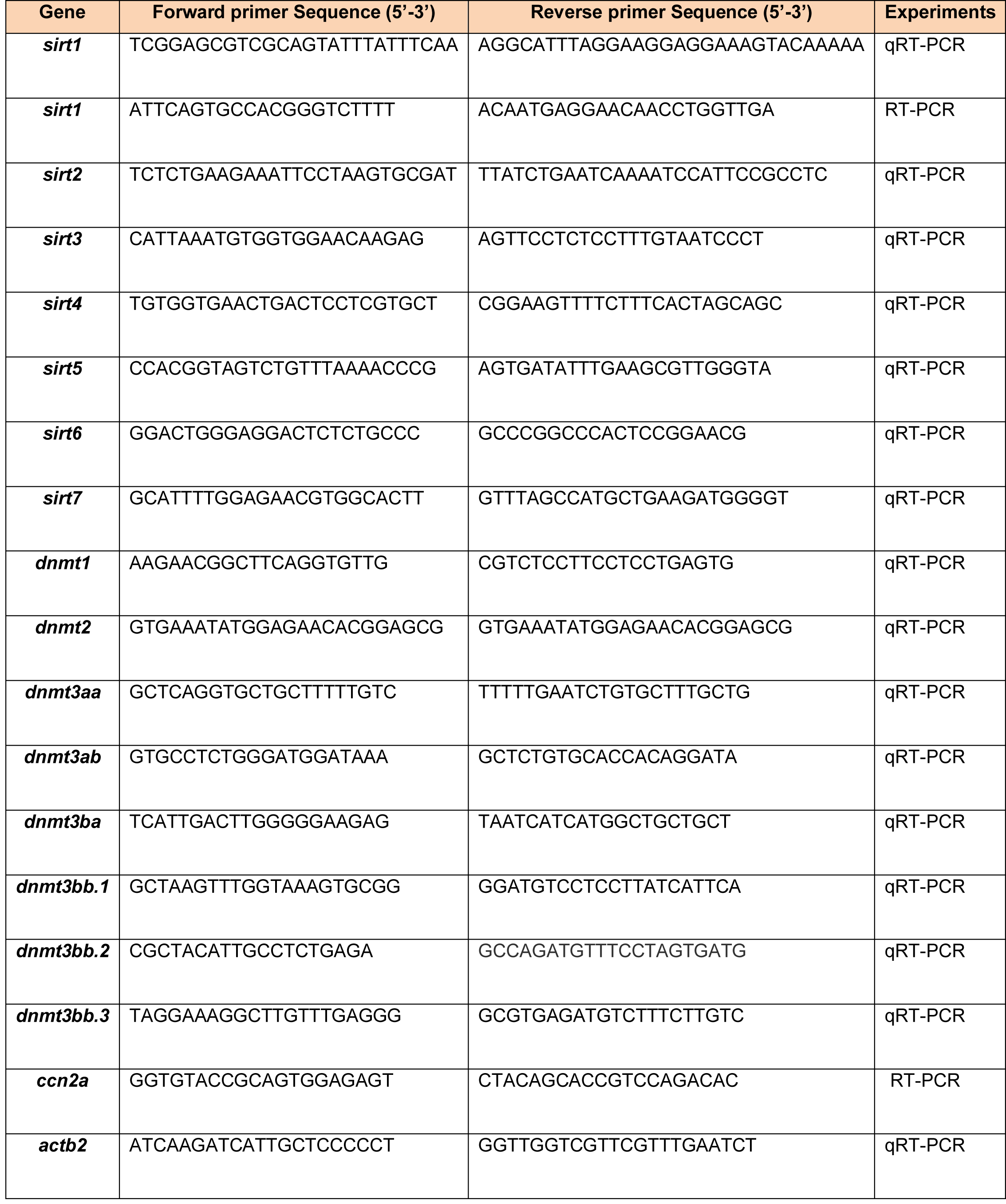
RT-PCR Primer list.

